# Lipoproteome screening of the Lyme disease agent identifies novel inhibitors of antibody-mediated complement killing

**DOI:** 10.1101/2021.09.23.461563

**Authors:** Michael J. Pereira, Beau Wager, Ryan J. Garrigues, Eva Gerlach, Joshua D. Quinn, Alex Dowdell, Marcia S. Osburne, Wolfram R. Zückert, Peter Kraiczy, Brandon L. Garcia, John M. Leong

## Abstract

Spirochetal pathogens such as the causative agent of Lyme disease, *Borrelia burgdorferi* sensu lato, encode an abundance of lipoproteins; however, due in part to their evolutionary distance from more well-studied bacteria such as Proteobacteria and Firmicutes, very few spirochetal lipoproteins have assigned functions. Indeed, *B. burgdorferi* devotes almost 8% of its genome to lipoprotein genes and interacts with its environment primarily through the production of at least eighty surface-exposed lipoproteins throughout its tick vector-vertebrate host lifecycle (57). Several *B. burgdorferi* lipoproteins have been shown to serve diverse roles, such as cellular adherence or immune evasion, but the functions for most *B. burgdorferi* surface lipoproteins remain unknown. In this study, we developed a *B. burgdorferi* lipoproteome screening platform utilizing intact spirochetes that enables the identification of previously unrecognized host interactions. As spirochetal survival in the bloodstream is essential for dissemination, we targeted our screen to C1, the first component of the classical (antibody-mediated) complement pathway. We identified two high-affinity C1 interactions by the paralogous lipoproteins, ErpB and ErpQ. Using biochemical, microbiological, and biophysical approaches, we demonstrated that ErpB and ErpQ inhibit the activated forms of the C1 proteases, C1r and C1s, and represent a new mechanistic class of C1 inhibitors that protect the spirochete from antibody-mediated complement killing by allosteric regulation. In addition to identifying a novel mode of complement inhibition, our study establishes a lipoproteome screening methodology as a discovery platform for identifying direct host-pathogen interactions that are central to the pathogenesis of spirochetes, such as the Lyme disease agent.

**Significance Statement:** Spirochetal pathogens encode an abundance of lipoproteins that can provide a critical interface with the host environment. For example, *Borrelia burgdorferi*, the model species for spirochetal biology, must survive an enzootic life cycle defined by fluctuations between vector (tick) and vertebrate host. While *B. burgdorferi* expresses over eighty surface lipoproteins— many of which likely contribute to host survival—the *B. burgdorferi* lipoproteome is poorly characterized. Here, we generated a platform to rapidly identify targets of *B. burgdorferi* surface lipoproteins and identified two orthologs that allosterically inhibit complement C1 subcomponents, conferring resistance to classical complement killing. This work expands our understanding of complement evasion mechanisms and points towards a discovery approach for identifying host-pathogen interactions that are central to spirochete pathogenesis.

## Introduction

The spirochete *Borrelia burgdorferi* sensu lato is the etiological agent of a diverse set of symptoms collectively referred to as Lyme disease, which is estimated to infect over 476,000 people annually in the U.S (1). *B. burgdorferi* is transmitted to humans and other reservoir hosts—primarily small mammals and birds—via the bite of a nymphal or adult-stage infected hard tick (*Ixodes scapularis)*. Upon tick feeding, bacteria are exposed to host blood in the tick midgut and then migrate to the salivary gland to be injected into the host dermis, where they establish a local spreading skin infection reflected in a characteristic expanding rash, erythema migrans (2, 3). The spirochetes then disseminate via the circulatory and/or lymphatic systems to colonize other sites, such as joints, heart, nervous tissue, and distant skin (4). Spirochetes can then be acquired by other feeding ticks, including larval stage ticks (5). As transovarial spread of *B. burgdorferi* does not occur in ticks, this feeding step is critical for intergenerational spirochetal transmission and retention of the bacterium in the tick population.

The ability of the spirochete to spread within the vertebrate host is reflected in its ability to cause multisystemic human disease, including arthritis, carditis, neuroborreliosis, and the formation of multiple erythema migrans lesions. The interaction of the Lyme disease spirochete with the host extracellular environment promotes its dissemination and persistence and is mediated, in part, by its surface lipoproteome. Spirochetal pathogens encode an abundance of lipoproteins, some of which are located on the bacterial surface (8, 69, 70), and in fact most of ∼125 *B. burgdorferi* lipoproteins are surface localized (6, 58). Many of these lipoproteins recognize identical or related host targets and/or interact with more than one host ligand (7). For example, at least 11 *B. burgdorferi* lipoproteins recognize host glycosaminoglycans (8), and nearly a dozen more interact directly with components of the innate immune system known as the complement cascade (9, 10). Understanding the interface between the complex *B. burgdorferi* surface lipoproteome and host macromolecules is fundamental to improving disease treatment and pursuing novel vaccine targets. However, due in part to their evolutionary distance from the better-studied bacteria such as Proteobacteria and Firmicutes, relatively few *B. burgdorferi* lipoproteins have assigned functions.

For both survival during exposure to the bloodmeal in the tick midgut and dissemination of the spirochete throughout the vertebrate host, protection against host defenses is essential. The complement system is the most immediate threat to survival that pathogens must contend with in the blood. This system is composed of a set of soluble and membrane-associated proteins that interact and activate a multistep proteolytic cascade upon detection of microbial surfaces, ultimately forming complexes that can damage microbial membrane integrity, recruit immune cells, and enhance phagocytosis (11–14). The three canonical pathways of complement system activation are each triggered by the recognition of molecular patterns on pathogenic surfaces. The lectin pathway (LP) proceeds by the recruitment of serine proteases (MASPs) to mannose-binding lectin (MBL) bound to the microbial surface by recognition of mannose or related sugars. The alternative pathway (AP) is triggered when complement factor C3 undergoes spontaneous self-cleavage in proximity of a microbial surface; it also serves as the central amplification loop of the complement cascade. The classical pathway (CP) typically initiates through the binding of host C1 to IgG or IgM complexes on the bacterial surface, although pathogen- or damage-associated molecular patterns can also trigger this pathway. All three pathways result in the formation of enzymatic complexes that trigger the release of proinflammatory peptides, the opsonization of the microbe, and the formation of a membrane attack complex (MAC) that lyses the pathogen.

To promote survival during tick feeding and/or spread within the vertebrate host, *B. burgdorferi* encodes surface lipoproteins that inhibit key steps of complement activation (9, 10, 71). *B. burgdorferi* OspC (Outer surface protein C), a lipoprotein essential to the spirochete life cycle, binds to C4b to inhibit *B. burgdorferi* bloodstream clearance (15). In addition, *B. burgdorferi* produces three distinct classes of Factor H binding proteins termed Complement Regulator Acquiring Surface Proteins (CRASPs), including CspA (CRASP-1), CspZ (CRASP-2), and ErpP/ErpC/ErpA (CRASP-3/CRASP-4/CRASP-5) (16–25). Each of these proteins binds Factor H, the major negative host regulator of the central amplification loop of the complement cascade and protects the bacterial surface from C3 deposition (59). The timing of expression varies among CRASPs, and CspA is specifically required for tick-to-host spirochete transmission, whereas CspZ mediates dissemination through the bloodstream and into distal tissues (26, 27).

Among known borrelial complement evasion proteins, *B. burgdorferi* BBK32 is unique in its ability to bind the complement C1 complex (28, 29). As the sole activator of the CP, C1 is comprised of the scaffold protein C1q and a heterotetramer of the serine proteases C1r and C1s (*i.e.,* C1qC1r_2_C1s_2_). C1q binding to the Fc region of an engaged antibody activates C1r to cleave C1s, which in turn cleaves complement components C2 and C4, leading to downstream C3 and C5 activation. BBK32 binds the C1 complex by recognizing C1r, blocking C1r proteolytic activity. When ectopically produced in a non-infectious, high-passage, otherwise serum-sensitive *B. burgdorferi* strain, BBK32 confers serum resistance (28). However, in an infectious strain background (*i.e.*, strain B31), a Δ*bbk32* mutant remains resistant to CP-mediated complement killing (28), suggesting that additional borrelial factors protect the spirochete from complement activation through this pathway.

*B. burgdorferi* carries as many as 21 endogenous plasmids, many of which are not stably maintained during *in vitro* culture, thus complicating genetic approaches to the identification of novel virulence factors (30). Nevertheless, a transposon library of *B. burgdorferi* has previously proved useful for genome-wide screens to identify many virulence factors (31). Unfortunately, functional redundancy of lipoproteins may limit its utility in exploring the genome for host interactions. Alternatively, gain-of-function studies have allowed researchers to detect the acquisition of new virulence-associated functions, such as complement resistance or cell attachment (28, 32, 72, 73). This is accomplished through ectopic lipoprotein production in a high-passage strain that, due to stochastic plasmid loss, lacks many virulence-associated functions and is non-infectious. To comprehensively identify *B. burgdorferi* lipoproteins located on the outer surface of the spirochete, Dowdell *et al.* ectopically produced epitope-tagged versions of all 127 putative lipoproteins encoded by *B. burgdorferi* strain B31 in the high passage strain B31-e2, finding that more than 80 are detected on the outer surface (6).

In this study, we used this library of B31-e2 clones to establish a surface lipoproteome screening methodology. Based on the serum resistance phenotype of a *bbk32-*deficient mutant described above and the observation that the complement evasion system of Lyme disease spirochetes has evolved to be functionally overlapping, we targeted our lipoproteome screen towards the human C1 complex. We found that two members of the Erp lipoprotein family, ErpB and ErpQ, bind C1 with high affinity and block its activity through a mechanism involving allosteric inhibition of the C1s protease subcomponent. Furthermore, we show that ErpB and ErpQ promote resistance to antibody-dependent complement killing. The discovery of a new role for ErpB and ErpQ in evading complement provides a validation of our lipoproteome screening methodology, which may be leveraged again in future studies to better understand the host-pathogen interface of the most prominent vector-borne pathogen in North America.

## Results

### Screening the *B. burgdorferi* surface lipoproteome identifies high-affinity interactions between ErpB and ErpQ with human C1

Utilizing a previously described lipoproteome library, we developed a whole-cell binding assay to screen 80 strains of *B. burgdorferi* B31-e2 that each ectopically overproduce a single distinct C-terminally His-tagged, surface-localized lipoprotein from the *B. burgdorferi* lipoproteome (6) for the ability to adhere to candidate ligands. As non-adherent controls, we included the parental strain B31-e2, as well as a strain that overproduces the periplasmic-localized lipoprotein BB0460. The 80 strains were previously shown to express surface-localized lipoproteins (6). To validate our approach, we first screened the library for strains that bind to human fibronectin. As expected, the two strains that bound fibronectin most strongly overexpressed the *B. burgdorferi* outer surface lipoproteins BBK32 and RevA, each of which have been shown to bind human fibronectin (34–38) (**Fig S1, Table S1)**.

To identify surface lipoproteins that target the classical complement pathway (CP), we screened the library for binding to purified, immobilized, human C1 complex. In addition to binding fibronectin and dermatan sulfate, BBK32 binds C1 (28, 29), and, as expected, spirochetes overexpressing BBK32 bound specifically to C1 in our screen (**Fig 1A, blue**). Interestingly, strains overexpressing lipoproteins ErpB or ErpQ also bound strongly to C1, exhibiting a relative signal higher than that of the BBK32-expressing strain (**Fig 1A**).

**Figure 1.**
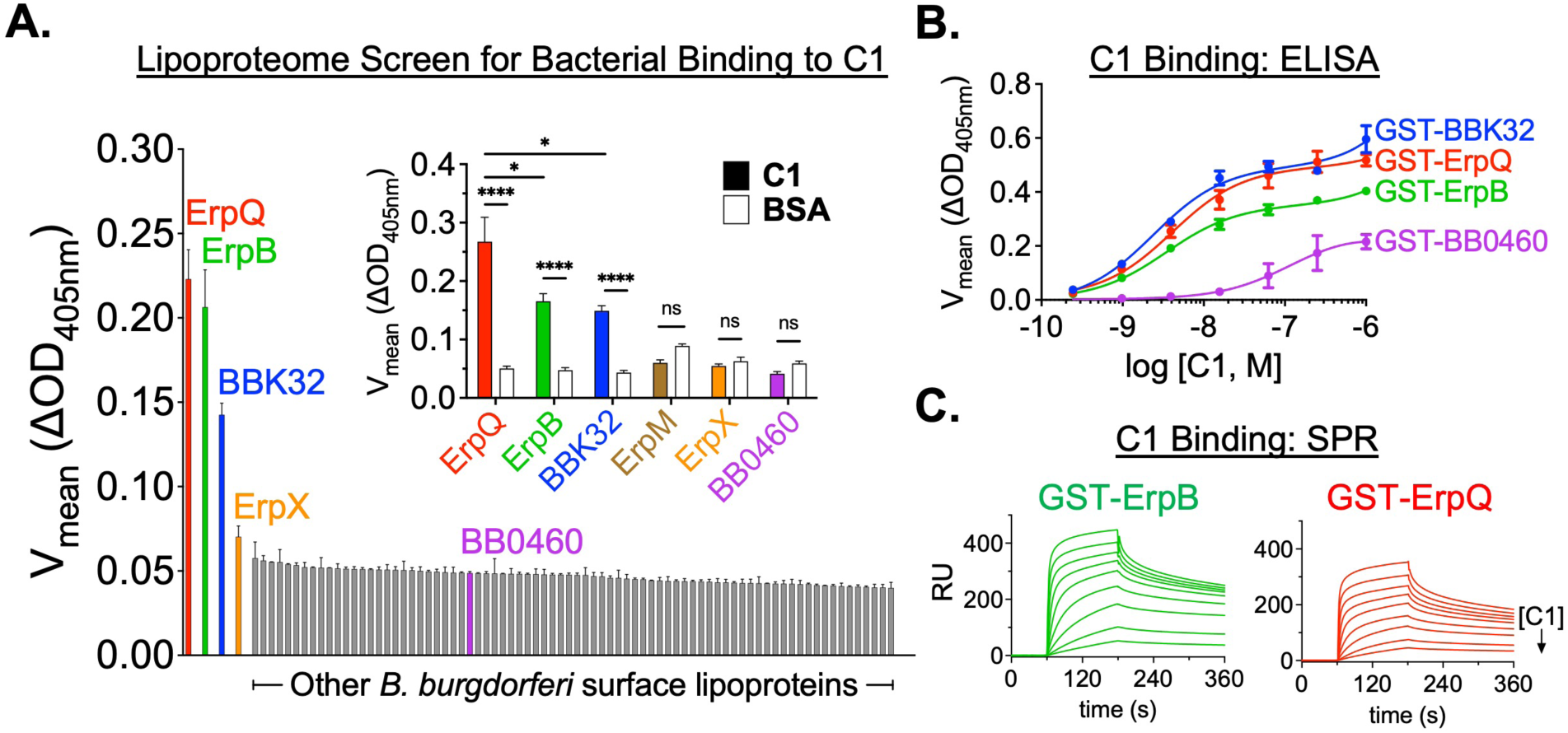
Screening the *B. burgdorferi* surface lipoproteome identifies high affinity interactions between ErpB and ErpQ with human C1. **(A)** 1×10^6^ strain B31-e2 producing one of 80 *B. burgdorferi* surface lipoproteins ((6); **Table S1**), as well as a periplasmic lipoprotein (BB0460) to serve as a negative control, were applied to microtiter wells coated with human C1 complex in duplicate. After washing, bound bacteria were quantitated by the change in OD_405nm_ over time by ELISA using an anti-*Bb* antibody (Abcam, ab20118). The clones are sorted in order of binding signal. Error bars indicate SEM. (**A, inset**) Binding of clones producing the indicated *B. burgdorferi* Elp protein, along with a positive control (BBK32) or a periplasmic negative control (BB0460), to immobilized human C1 complex or BSA was quantitated as described above. Error bars indicate SEM. ****, p<0.0001; *, p<0.05; ns, not significant using Student’s *t* test to compare mean values. (**B)** Binding of the indicated GST-fusion proteins to wells coated with the indicated concentration of human C1 complex was quantitated. The experiment was performed six times (GST-ErpB) or nine times (GST-BBK32 and GST-ErpQ) at each concentration and error bars indicate SEM. Affinity analysis was performed with Prism GraphPad software, using a non-linear regression analysis. (**C)** The ability for GST-ErpB (left) GST-ErpQ (right) to bind human C1 complex was evaluated by SPR. A two-fold dilution series (0.6 - 150 nM) of C1 complex was injected over GST-ErpB and GST-ErpQ biosensors and steady-state affinity analysis was carried out with T200 Evaluation Software. Each SPR experiment was performed in triplicate. Equilibrium dissociation constants (*K*_D_) calculated from ELISA-type and SPR binding assays are shown in **Table 1**.

**Table 1.**
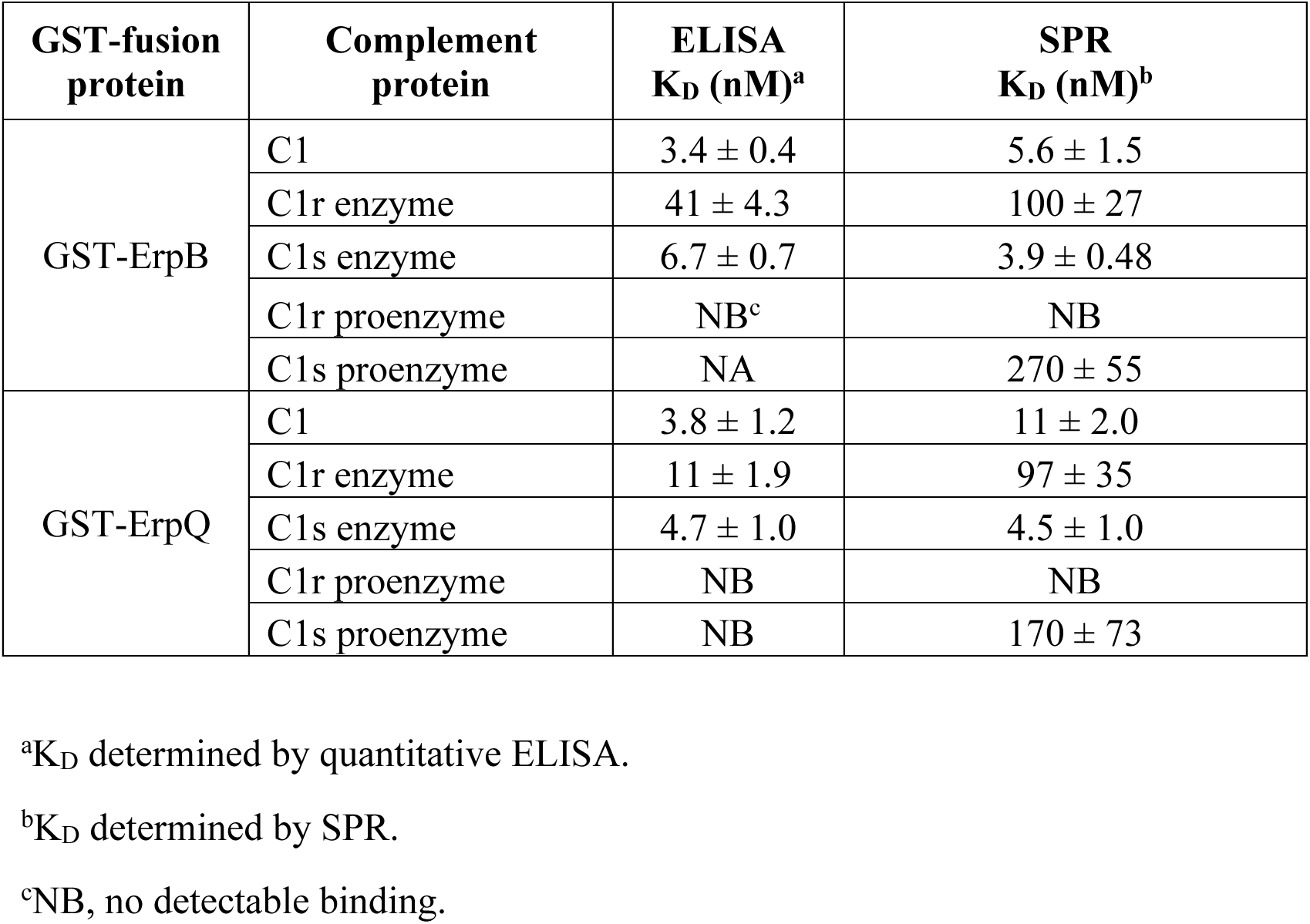

ErpB and ErpQ are members of the *B. burgdorferi* OspEF-related protein family (Erps) (39–41). All *erp* genes are encoded on circular plasmid 32 DNA elements (cp32), and in *B. burgdorferi* strain B31, ten cp32 plasmids together encode 13 Erp proteins (39–41). Of these, five belong to the Elp subfamily of Erps, which includes ErpB and ErpQ, and is defined by OspE/F-like leader peptides (Elps) (42). In addition to ErpB and ErpQ, the *B. burgdorferi* strain B31 genome includes Elp members ErpM, ErpO, and ErpX (**Table S2**). Despite being encoded on separate cp32 plasmids, *erpB* and *erpO* are identical at the amino acid sequence level, and for simplicity, ErpO will be referred to as ErpB hereafter. In strain B31, the Elp proteins (i.e., ErpB, ErpM, ErpQ, and ErpX) are 44-59% identical and 59-76% similar and exhibit their highest identity in the N-terminal and C-terminal protein regions (**Fig S2, Table S2**).

To confirm the results of our screen, and because little is known about the function of Elp proteins, we individually tested strains producing each Elp in the ELISA-based spirochete binding assay against the C1 complex, including bovine serum albumin (BSA) as a negative control (**Fig 1A, inset**). Spirochetes expressing BBK32 (a C1-binding protein) and BB0460 (a periplasmic-localized lipoprotein (6)), were used as positive and negative controls, respectively. Strains producing ErpB, ErpQ, or BBK32 all exhibited statistically significant binding to C1 relative to BSA, whereas ErpM, ErpX, and BB0460 did not (**Fig 1A, inset**).

To further investigate the ability of ErpB and ErpQ to directly bind to human C1, we purified recombinant GST-tagged fusion proteins (GST-ErpB and GST-ErpQ). Consistent with data obtained from the spirochete binding assay (**Fig 1A**), GST-ErpB and GST-ErpQ bound with high-affinity to immobilized C1 in an ELISA-type binding assay, exhibiting apparent equilibrium dissociation constants (*K*_D_) of 3.4 nM and 3.8 nM, respectively (**Fig 1B, Table 1**). To gain insight into the interaction of ErpB and ErpQ with soluble C1, we used surface plasmon resonance (SPR) whereby GST-ErpB and GST-ErpQ were immobilized on SPR sensor chips. When C1 was used as an analyte, strong C1-binding was observed, with GST-ErpB and GST-ErpQ exhibiting steady-state calculated *K*_D_ values of 5.6 and 11 nM, respectively (**Fig 1C, Table 1**). Together, these data confirm that ErpB and ErpQ individually promote spirochete binding to human C1 via direct interaction with this molecule.

### ErpB and ErpQ selectively bind the activated forms of C1r and C1s

The C1 complex is composed of C1q and a heterotetramer of C1r and C1s (i.e. C1r_2_C1s_2_) (**Fig S3A**). C1q is a non-enzymatic component and functions in pattern recognition, while C1r and C1s are serine proteases that catalyze the initial proteolytic reactions of the CP. To clarify whether ErpB and ErpQ bind to C1 by interacting with individual subcomponents, we carried out an ELISA-type binding assay using purified immobilized C1q and activated forms of C1r and C1s (i.e. C1r enzyme and C1s enzyme). Relative to the negative control GST-BB0460, no significant interaction was detected for either GST-ErpB or GST-ErpQ with human C1q, (**Fig S3B**). In contrast, each protein bound with high affinity to C1r enzyme (*K*_D_ of GST-ErpB/C1r = 41 nM; GST-ErpQ/C1r = 11 nM) as well as to C1s enzyme (*K*_D_ of GST-ErpB/C1s = 6.7 nM; GST-ErpQ/C1s = 4.7 nM) (**Fig S3C, D, Table 1**).

To further study the interaction of C1r and C1s with ErpB and ErpQ, we utilized Far-western blot analysis, detecting the His tag on the ectopically produced lipoprotein. We.first assessed the apparent molecular weights of ErpB and ErpQ in bacterial lysates by conventional western blotting, Pronase treatment was used to assess the surface-localization of each protein, as previously described (6). As expected, ErpB and ErpQ were predominantly expressed on the spirochetal surface and were detected as bands migrating at 61 kDa and 55 kDa, respectively. Full-length ErpQ was produced at higher levels than ErpB, and the presence of a prominent lower molecular weight ErpB band—presumably a stable degradation product—suggested that ErpB, but not ErpQ was subjected to proteolytic cleavage (**Fig S4A-C**). The higher level of ErpQ production correlated with a somewhat shorter bacterial length when observed under darkfield microscopy. We then probed these bacterial lysates using purified human C1 or the C1 subcomponent proteases to test for potential protein-protein interactions. Lysates from spirochetes expressing BBK32 (a C1r-binding positive control) contained a species that bound strongly to C1 complex, C1r proenzyme, and C1r enzyme, but, as expected, to neither form of C1s (**Fig 2 A, B**). In all cases the C1/C1r-binding species correlated with epitope-tagged BBK32 (**Fig S4A**). The negative control BB0460 lysates contained no species that bound detectably to any complement protein probe (**Fig 2 A, B**). Consistent with the data shown in **Figs 1 and S3**, single bands coincident with ErpB and ErpQ, as judged by an α-6xHis blot (**Fig S4A**), bound to C1 complex, C1r enzyme, and C1s enzyme (**Fig 2 A, B**). Furthermore, this binding was reduced in the lysates of cells treated with pronase (**Fig 2 A, B**).

**Figure 2.**
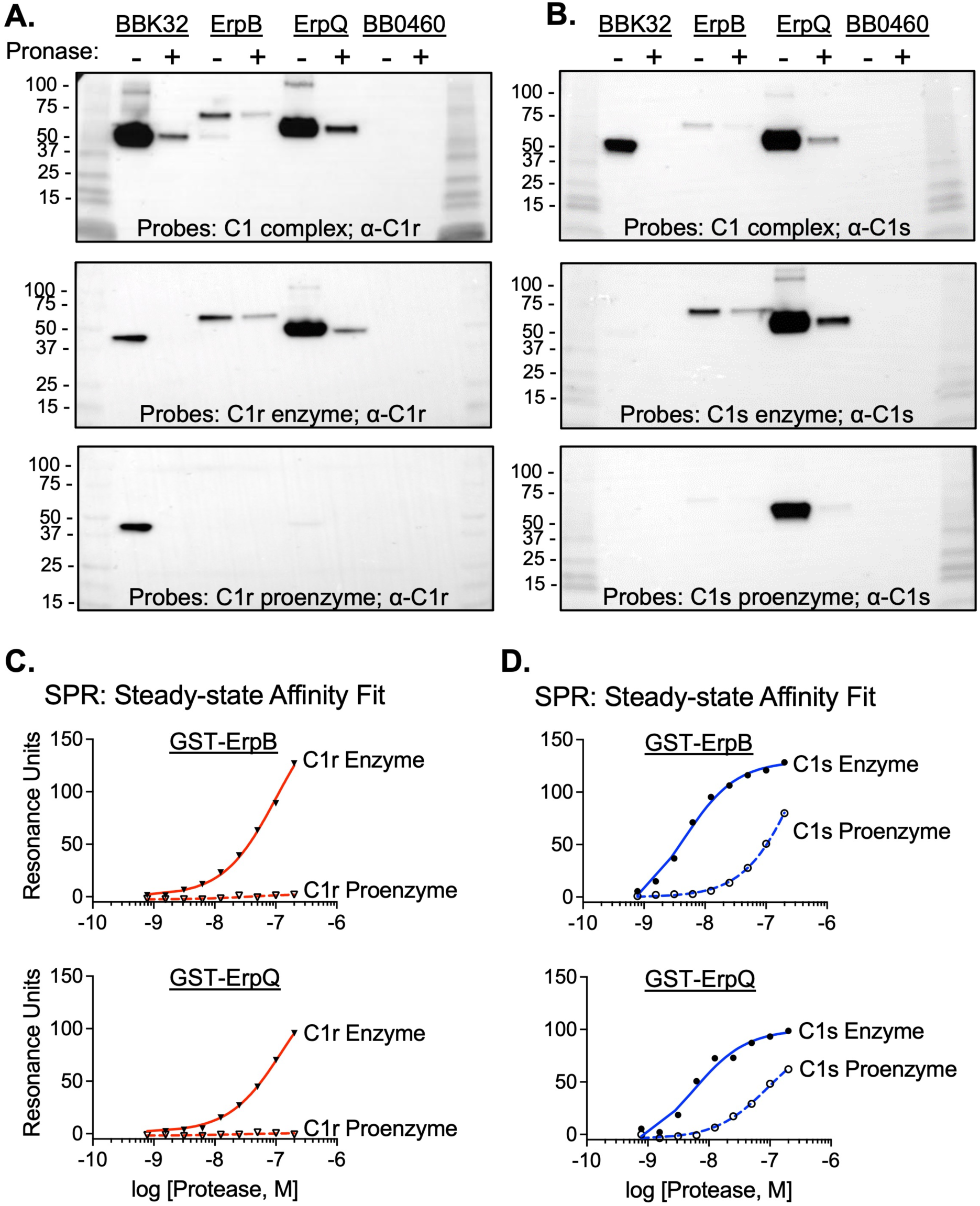
ErpB and ErpQ preferentially bind activated forms of C1r and C1s. **A)** Extracts from untreated (“-“) or pronase-treated (“+”) 1×10^7^ strain B31-e2 spirochetes that ectopically produce the indicated surface lipoproteins were separated by SDS-PAGE and transferred to PVDF membranes. The filters were probed with purified C1 complex (top), C1r enzyme (middle) or C1r proenzyme (bottom), and bound probe revealed by anti-C1r antibody, followed by HRP-conjugated anti-mouse antibody. Shown is a representative of 3 experiments. **B)** Filters prepared identically to panel A were probed with purified C1 complex (top), C1s enzyme (middle) or C1s proenzyme (bottom), and bound probe revealed by anti-C1s antibody, followed by HRP-conjugated anti-mouse antibody. Shown is a representative of 3 experiments. **C and D)** Biosensors immobilized with GST-ErpB (top) or GST-ErpQ (bottom) were tested by SPR for binding to the indicated concentrations of the enzyme or proenzyme forms of C1r **(C)** or C1s **(D)** Injection series were each performed in triplicate. For both panels C) and D), steady-state affinity fits were determined by T200 Biacore Evaluation software and *K*_D_ values are reported in **Table 1**.

Interestingly, we found that C1r proenzyme failed to bind either ErpB or ErpQ spirochete lysates (**Fig 2A**). Similarly, C1s proenzyme showed lower relative binding to ErpB and ErpQ compared to the activated form of C1s (**Fig 2B**). To follow up on this intriguing finding, we measured the relative affinities of pro- and active forms of both C1r and C1s for recombinant GST-ErpB and GST-ErpQ by SPR. Indeed, while GST-ErpB and GST-ErpQ bound to C1r enzyme with *K*_D_ values of 100 nM and 97 nM, respectively, neither protein exhibited detectable binding for C1r proenzyme (**Fig 2C, S5**). Similarly, GST-ErpB and GST-ErpQ bound C1s enzyme with ∼70-fold and ∼38-fold higher affinity, respectively, than C1s proenzyme (*K*_D_ = 3.9 nM vs. 270 nM; *K*_D_ = 4.5 nM vs. 170 nM) (**Fig 2D**, **S5, Table 1**).

### ErpQ inhibits C1s cleavage of C2 and C4

Having established that ErpB and ErpQ were capable of direct interaction with human C1 via specific recognition of the protease subcomponents, using ErpQ we explored a potential mechanism of action for C1 inhibition. To facilitate clarity in our gel-based cleavage assays and to eliminate the GST-tag from the mechanistic analysis, we generated an ErpQ construct lacking this epitope. The “tagless” ErpQ behaved nearly identically in SPR C1s-binding assays and ELISA-based complement assays when compared to GST-ErpQ (**Fig S6**).

Previously we have shown that BBK32, which binds to C1r but not C1s, is capable of directly inhibiting purified C1r enzyme cleavage of C1s proenzyme (28). In contrast, recombinant ErpQ failed to block this reaction at protein concentrations several orders of magnitude greater than the C1r/ErpQ K_D_ (**Fig S7A**). ErpQ also failed to prevent the cleavage of the small peptidic C1r substrate Z-Gly-Arg-sBzl (60), whereas BBK32 did so readily (**Fig S7B**). Similarly, unlike futhan, the small molecule active site C1s inhibitor (60), 25 µM ErpQ (i.e., > 5,500 fold over the measured *K*_D_, **Table 1**) failed to inhibit the cleavage of the C1s peptidic substrate Z-L-Lys thiobenzyl by C1s, (**Fig 3A**). Thus, in the C1s/ErpQ complex, the active site of C1s remains accessible to a small peptide substrate.

**Figure 3.**
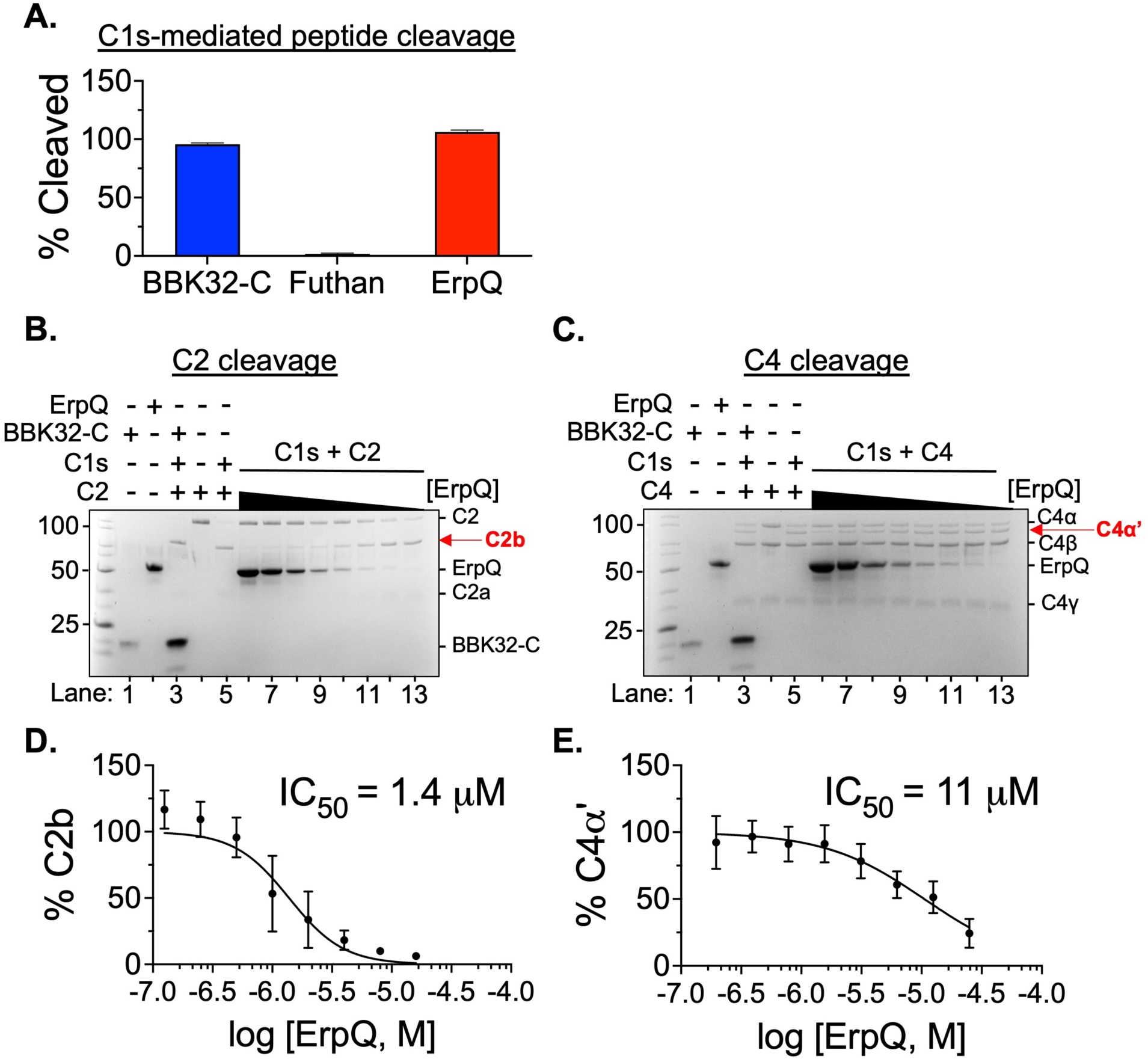
ErpQ is an allosteric inhibitor of complement C1s. **A)** Enzymatic cleavage by C1s of the small peptide substrate Z-L-Lys-sBzl was assayed with DTNB (Ellman’s reagent) in the presence of 25 μM BBK32-C (non-inhibitory control) or ErpQ at 25°C for 1hr. Experiments were performed in triplicate. Absorbance was read at 412 nm and signals were normalized to negative control no-substrate wells. **B)** Top: Proteolytic cleavage of C2 by C1s enzyme produces ∼70kDa C2b and ∼35kDa C2a after 1hr at 37°C. Lanes 1-5: C2b accumulation in the presence (“+”) or absence (“-“) or 25 µM ErpQ, 25 µ M BBK32-C (non-inhibitory control), 6.25 nM C1s, and 685 nM C2. (Note that the amount of C1s loaded is below the level of detection by SDS-PAGE). Lanes 6-13: C2b accumulation in the presence of 6.25 nM C1s, 685 nM C2 and a two-fold dilution series (from 16 to 0.13 μM) of ErpQ. Bottom: The fraction of C2b relative to total input C2 in the same lane determined by densitometry analysis data are normalized to C2 (lane 5) and C1s digested C2 (lane 6). A representative gel is shown. The experiment was performed three times. **C)** Top: C4, which consists of 3 polypeptide chains, C4α (97 kDa), C4β (77 kDa), C4γ (33 kDa), is cleaved by C1s enzyme for 1hr at 37 °C to produce C4α’ (88 kDa). Lanes 1-5: SDS-PAGE profile in the presence (“+”) or absence (“-“) or 25 µ M ErpQ, 25 µ M BBK32-C (non-inhibitory control), 6.25 nM C1s, and 616 nM C4. Lanes 6-13: SDS-PAGE profile in the presence of 6.25 nM C1s, 616 nM C4 and a two-fold dilution series (from 25 to 0.20 μM) of ErpQ. Bottom: The fraction of C4α’ relative to input C4β in the same lane and normalized to C1s + C4 positive control (lane 6) and negative control C4 (lane 5) was determined by densitometry analysis.

We next tested whether ErpQ was capable of inhibiting C1s-mediated cleavage of native substrates. The cleavage of C2 or C4 by purified C1s was monitored by SDS-PAGE in the presence of increasing concentrations of ErpQ (**Fig 3B,C**). Whereas BBK32 failed to block C2 cleavage by C1s (**Fig 3B, lane 3**) to generate the cleavage product C2b (“ß **C2b**”; **Fig 3B**), ErpQ blocked C1s-mediated C2 proteolysis and the concomitant formation of C2b in a dose-dependent fashion (**Fig. 3B, lanes 6-13**). Likewise, while BBK32 failed to prevent C4 cleavage by C1s (**Fig 3C, lane 3**) to generate the cleavage product C4α’ (“ß **C4α’**”; **Fig 3C**), ErpQ did so in a dose-dependent manner (**Fig. 3C, lanes 6-13**). Densitometry analysis resulted in calculated ErpQ IC_50_’s of 1.4 µM and 11 µM for C2 and C4, respectively. The observation that ErpQ inhibited the cleavage of large endogenous C1s substrates but not a small peptide C1s substrate suggests that ErpQ inhibits C1s allosterically, leaving the active site of C1s accessible to small peptides.

### ErpB and ErpQ inhibit the classical pathway of complement

Collectively, the data above identify a novel interaction between surface-expressed *B. burgdorferi* lipoproteins ErpB and ErpQ with human C1 and demonstrate that recombinant ErpQ blocks C1s activity. The CP is initiated by this C1 activity, so we tested the ability of ErpB and ErpQ to block successive steps in this pathway. Recombinant GST-ErpB or GST-ErpQ fusion proteins were added at increasing concentrations to normal human serum in microtiter wells coated with IgM to initiate CP activation. The surface deposition of C4b, C3b, and C5b-9, mimicking the fixation of successive components of the CP (59) was measured by ELISA. GST-BBK32 and GST-BB0460 served as positive and negative controls, respectively. Both GST-ErpB and GST-ErpQ inhibited the deposition of these three components in a dose-dependent manner, with half-maximal inhibitory concentrations (IC_50_’s) approximately ten-fold higher than the IC_50_ of GST-BBK32 (**Fig 4A-C, Table 2**). GST-BB0460 showed no inhibitory activity. As C5b-9 is the membrane attack complex, capable of generating pores in membranes, we further tested each protein for protection of antibody-sensitized sheep red blood cells from CP-mediated lysis. As above, GST-ErpQ and GST-ErpB inhibited lysis in a dose-dependent manner, with an IC_50_ of 1.5 and 1.6 μM, respectively, or ∼20-fold higher than the IC_50_ of GST-BBK32 **(Fig 4D, Table 2**).

**Figure 4.**
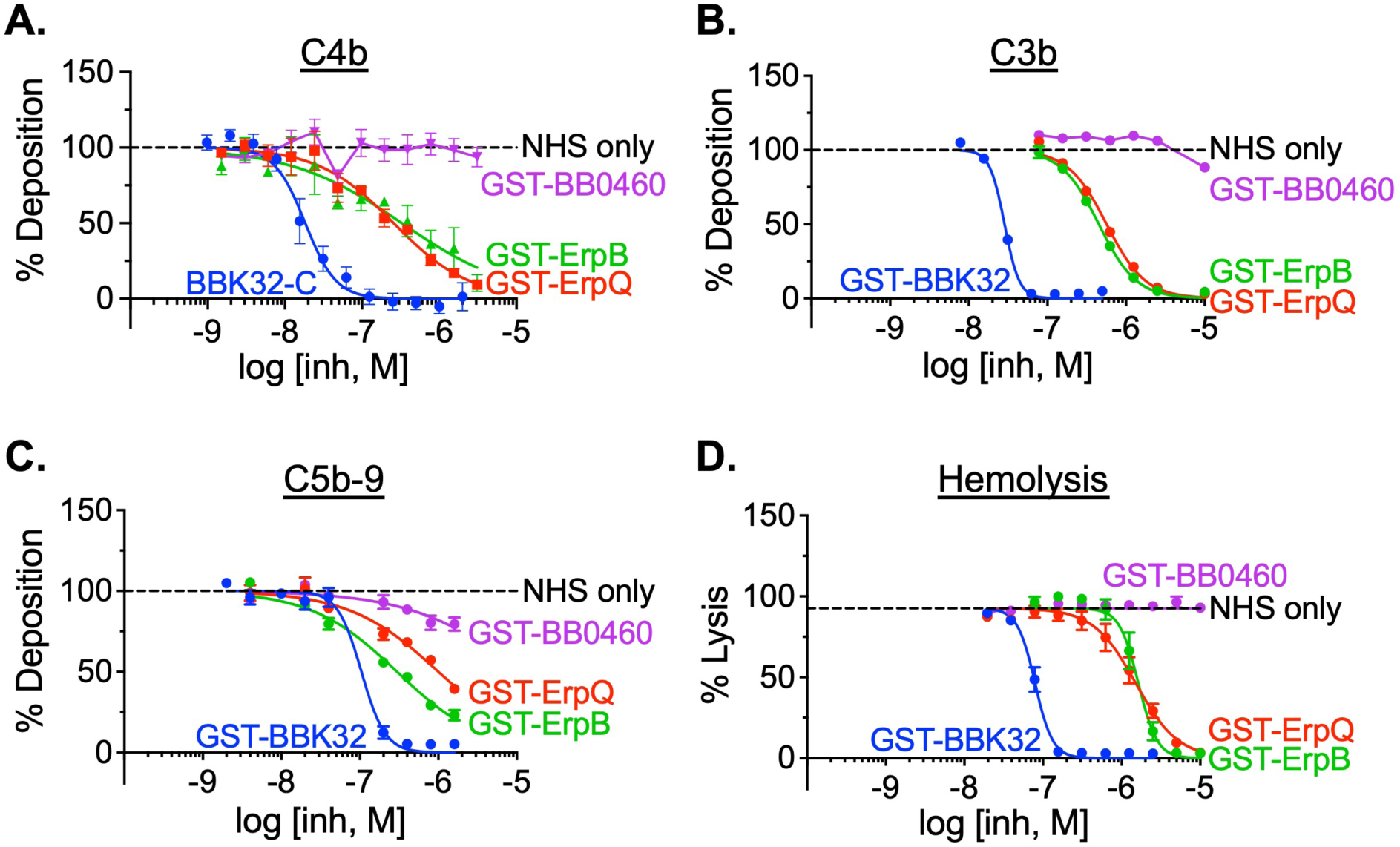
ErpB and ErpQ inhibit the classical pathway of complement. **A-C)** Normal human serum (NHS) was incubated with the indicated concentration of purified GST-fusion proteins, then added to wells precoated with human IgM. Deposition of **A)** C4b, **B)** C3b, or **C)** C5b-C9 was determined by the addition of the appropriate primary and secondary antibodies (see Materials and Methods) enumerated by absorbance at OD_405nm_ or OD_450nm_. Each well was normalized to wells with no inhibitor (100%) and no serum (0%). Curves were fit using nonlinear regression to determine IC_50_ values. **D)** NHS was incubated with the indicated concentration of purified GST-fusion proteins and then added to pre-opsonized sheep erythrocytes (see Materials and Methods). Erythrocyte lysis was determined by OD_405nm_ and normalized to lysis by deionized water (100%) and no serum (0%). Error bars indicate SEM. Each concentration was tested a minimum of three times.

**Table 2.** IC_50_ for ErpB- and ErpQ-mediated inhibition of the classical pathway of complement.

| **Protein** | **Hemolysis** | **C5b-9** | **C3b** | **C4b** |
| --- | --- | --- | --- | --- |
| GST-BBK32 | 79 ± 5.2 | 100 ± 20 | 29 ± 1.0 | 18 ± 3.3* |
| GST-ErpQ | 1500 ± 240 | 1000 ± 210 | 450 ± 26 | 260 ± 62 |
| GST-ErpB | 1600 ± 190 | 300 ± 43 | 570 ± 29 | 330 ± 130 |

\*IC<sub>50</sub> based on non-GST-tagged BBK32-C construct.

### Ectopic production of ErpB and ErpQ protect spirochetes from complement-mediated killing

The ability of recombinant GST-ErpB and GST-ErpQ to block complement deposition products and prevent lysis of red blood cells by the membrane attack complex suggested that these proteins may protect spirochetes from antibody-dependent complement attack. We tested the ability of *B. burgdorferi* B31-e2 strains that ectopically produce (His-tagged) ErpB or ErpQ (Fig 1A) to resist CP killing, with BBK32 and BB0460 as positive and negative controls, respectively. Based on a previously-described assay to initiate the CP (61), we incubated these strains with *B. burgdorferi*-specific polyclonal antibodies, then added normal human serum to provide complement components and lysozyme to facilitate disruption of spirochetal integrity (see Methods). After dilution into BSK-II media and 72-hour incubation to allow for growth of surviving bacteria, we enumerated living spirochetes. Treatment with isotype control antibody or heat-inactivated serum were utilized as negative controls, and survival indices were calculated after normalization to spirochetes surviving treatment with heat-inactivated serum.

As predicted, a *B. burgdorferi* B31-e2 high-passage strain that ectopically produced the periplasmic protein BB0460 was highly susceptible to antibody-dependent complement killing, with an index of less than 0.3% (**Fig 5E, purple**). Conversely, production of BBK32 conferred high-level protection, with an index of ∼56%, or ∼190-fold higher than the negative control BB0460 (**Fig 5E, blue**). Spirochetes producing ErpB or ErpQ displayed survival indices of 9- and 56-fold higher, respectively, than the those producing BB0460 (**Fig 5E, green and red**). Compared to *B. burgdorferi* B31-e2 producing BB0460, the strain producing ErpQ displayed a six-fold defect in survival index after incubation with isotype control antibody, suggesting that the overproduction of ErpQ may moderately enhance susceptibility of the spirochete to non-CP serum killing. Nevertheless, the dramatically enhanced resistance to CP-mediated killing conferred by ErpB and ErpQ indicates that the inhibition of C1s cleavage and inactivation of the CP observed in biochemical assays reflects an activity that protects bacterial viability.

**Figure 5.**
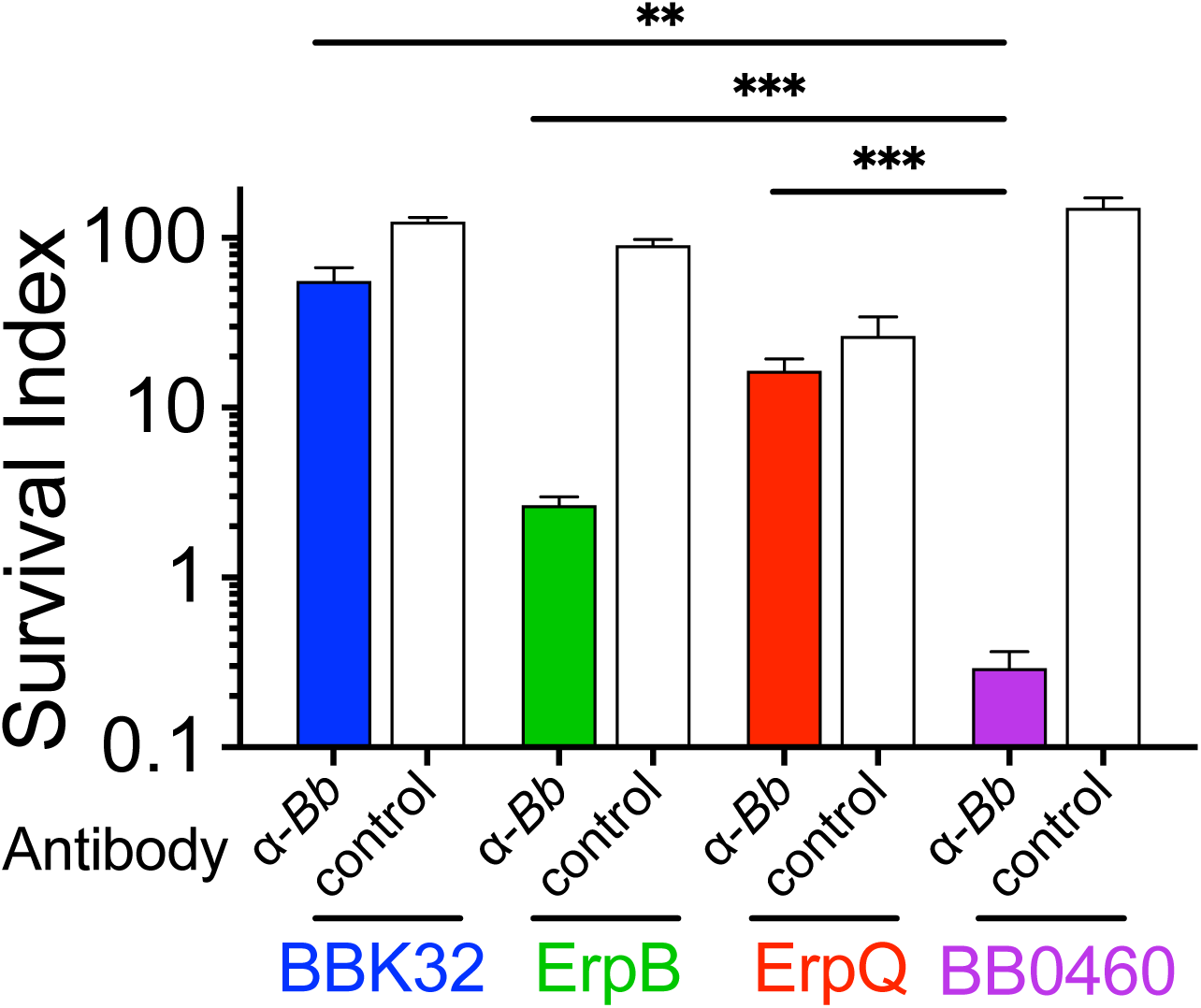
Ectopic production of ErpB and ErpQ protect spirochetes from complement-mediated killing. 5×10^7^ spirochetes were treated with a 4 μg/ml of an α-*B. burgdorferi* strain B31 mouse polyclonal antibody or its isotype (IgG) control, followed by exposure to 20% untreated or heat-inactivated NHS (or heat-inactivated NHS) containing 10 μg/ml lysozyme for four hours. Samples were grown in BSK-II for 72 hours at 33°C and enumerated by dark field microscopy. Survival index was calculated by dividing the culture density of each sample in the “Bb-specific Ab” or “Isotype control Ab” groups by the average density of the heat-inactivated serum group for each strain. Shown is the mean and SEM of triplicate samples. ***, p<0.001; **, p<0.01; ns, not significant using Student’s *t* test to compare mean values.

## Discussion

Lyme disease spirochetes are typical of other spirochetal pathogens in that they encode many lipoproteins (58). Although the proportion of lipoproteins located in the periplasm varies among spirochetes (6, 69, 74, 75), surface lipoproteins are critical to pathogenesis and provide an important means by which pathogenic spirochetes interact with the host environment and (76, 77). Of the approximately125 lipoproteins encoded by the *B. burgdorferi* genome, the majority localize to the outer membrane (6), although functions for relatively few of these proteins have been elucidated. Adding to the complexity of understanding lipoprotein function, several of the best characterized *B. burgdorferi* outer surface lipoproteins, such as OspC and BBK32, have been shown to provide multiple independent functions during murine borreliosis (43–47). Building on the generation of a comprehensive lipoprotein library (6), we developed a screening methodology to identify novel interactions between host macromolecules and the *B. burgdorferi* surface lipoproteome expressed in its native environment in the outer membrane of intact spirochetes. This methodology has the potential to uncover diverse host interactions that take place at the spirochete surface and may be valuable in the study of other pathogenic bacteria as well.

As an extracellular pathogen that encounters host blood during both the tick bloodmeal and throughout dissemination and colonization of their vertebrate hosts, Lyme disease spirochetes must prevent complement-mediated opsonization and lysis at multiple stages in the enzootic cycle. Moreover, the complement system employs three distinct pathways for activation that together form a complex host defense. Reflecting this, nearly a dozen different *B. burgdorferi* outer surface lipoproteins have been shown to directly interact with complement components, disrupting their activities (9, 10). At least three factors contribute to the multiplicity of lipoproteins devoted to thwart complement defense. First, distinct borrelial complement evasion proteins block different complement activation pathways. For example, BBK32 selectively targets C1r, the initiator protease of the classical pathway, while OspC binds to C4b, the downstream activation product of both the classical and lectin pathways (15, 28, 29). Second, individual borrelial lipoproteins may target the same host protein but function at different stages of the enzootic cycle. *B. burgdorferi* CspA and CspZ both bind to factor H and prevent activation of the alternative pathway, but CspA is expressed exclusively in the tick midgut and prevents the bacteriocidal effects of the bloodmeal, whereas CspZ is produced early in vertebrate infection and fosters the establishment of infection in that host ( 26, 27, 78). Finally, although some Lyme disease spirochete strains are restricted to only a single vertebrate, other strains have the capacity to infect multiple vertebrate hosts (62) that encode polymorphic complement components (63, 64). Indeed, variation in CspA sequences have been shown to dictate binding to mammalian vs. avian factor H and the concomitant capacity to infect these two hosts (27, 66). Likewise, the production of multiple complement-inactivating proteins may permit the broad host specificity displayed by some *B. burgdorferi* strains. Thus, the collective activities of multiple complement evasion proteins of *B. burgdorferi* may provide the distinct temporal and spatial needs to thrive in enzootic cycles that involve multiple hosts. Due to the complexity of these interactions. *B. burgdorferi* serves as a useful model for understanding how a wide range of complement inactivation mechanisms together foster the retention of a pathogen in nature.

Consistent with the observation that partial functional redundancy is a hallmark of the *B. burgdorferi* complement evasion system, BBK32 was sufficient to protect spirochetes from complement-mediated killing, but *bbk32*-deficient mutants remained serum resistant (28). Thus, we focused our surface lipoproteome screen on the classical pathway component C1. We identified two members of the paralogous Elp protein family, ErpB and ErpQ (from *B. burgdorferi* strain B31), as capable of forming high affinity interactions with the human C1 complex (**Figs 1, 2, S3**); both proteins are antigenic during experimental murine and human infection, indicating that they are produced *in vivo* (83, 84, 85). The Erp family encompasses more than 17 genes in strain B31 (86) that share highly homologous leader peptides and DNA sequence at the 5’ end of their operons (39, 42, 83). However, the amino acid sequences of their mature proteins group them into the OspE-related, OspF-related, and Elp subfamilies that are evolutionarily unrelated (42). Many OspE-related proteins have been shown to bind factor H (87, 88, 89, 90, 91), and several OspF-related proteins bind to heparan sulfate (32). Our finding that two Elp members bind to complement C1 further supports the hypothesis of divergent functions among the three subfamilies (42).

Consistent with the mechanistic divergence of anti-complement lipoproteins, ErpB and ErpQ, like BBK32, prevent antibody-mediated complement activation but target the C1 complex via distinct means. BBK32 does not bind C1s, but recognizes both zymogen and activated forms of C1r, blocking its enzymatic activity. In contrast, ErpB and ErpQ bind to both C1r and C1s but selectively recognize activated forms of the proteases (**Figs 2, S5**). Further, we showed that ErpQ in incapable of blocking C1r activity (**Fig S7**) but prevents cleavage of both C2 and C4 by activated C1s enzyme (**Fig 3B)**; it seems highly likely that ErpB possesses a similar activity. Finally, ErpQ did not prevent cleavage of small peptide substrates, and is unusual among microbial-derived serine protease inhibitors, such as ecotin or BBK32 (49), which typically target the active site (79, 80).

Previous work showed that expression of BBK32 by a high-passage, noninfectious *B. burgdorferi* strain enhanced serum resistance, and that simultaneous inactivation of the classical and lectin pathways eliminated this enhancement, indicating that BBK32 blocked one or both pathways. To confirm that the C1-binding activities of BBK32, ErpB and ErpQ specifically blocked classical complement killing, we triggered this pathway by treating high-passage strains that ectopically produce these proteins with anti*-B. burgdorferi* antibody. Whereas BBK32, ErpB and ErpQ provided no survival advantage when spirochetes were treated with serum supplemented with isotype control antibody, all three lipoproteins promoted survival when incubated with specific antibody, indicating that the C1r- or C1s-inhibitory activities of BBK32 or ErpB and ErpQ, respectively, protected spirochetes from classical complement killing. BBK32 provided the greatest degree of protection, enhancing the survival index 190-fold relative to BB0460, compared to 56-fold and 9-fold for ErpQ and ErpB, respectively, (**Fig 5)**. Notably, BBK32 and ErpQ appeared to be expressed at much higher levels than ErpB (**Fig 2A**). In addition, BBK32 inhibited C4b and C3b in vitro deposition and complement-mediated RBC hemolysis at ∼10-fold lower concentrations than ErpB or ErpQ (**Fig 4**).

The innate and adaptive immune system intersect at the level of the classical pathway of complement when antibody-antigen immune complexes are recognized by complement C1, triggering the complement cascade. Blocking complement C1 may be critical for *B. burgdorferi* persistence in immunocompetent hosts, which generate a specific antibody response during chronic infection. This activity might also be required to establish infection in a previously infected host, or, given that natural antibodies recognize the Lyme disease spirochete (50), in a naïve host.

ErpB and ErpQ display identical biochemical activities, and no evidence to-date has indicated divergent expression patterns between the two genes, raising the possibility that they are functionally redundant. In addition, other Elp family members such as ErpX and ErpM (which is as closely related to ErpB and ErpQ as they are to each other; Table S3) did not bind human C1 (**Fig 1A**). Complement C1 is polymorphic among vertebrates, and whether these Elp’s recognize C1 of other *B. burgdorferi* hosts, perhaps contributing to host specificity, remains to be tested (81). Historically, comprehensive analysis of *B. burgdorferi* gene families has been limited by the difficulty of genetic manipulation of infectious strains. However, recent adaptation of CRISPRi to this pathogen (82) may enable future comprehensive examinations of the role of Elp proteins during the enzootic lifecycle of *B. burgdorferi*.

## Materials and Methods

### Expression plasmid cloning and protein purification

All primers used in this study are listed in TableS4. To generate expression plasmids encoding N-terminal GST fusions of ErpB, ErpQ, BBK32, and BB0460, genomic DNA was first prepared from these *B. burgdorferi* B31-e2 expression strains using the DNeasy Blood and Tissue Kit (Qiagen). The open reading frame (lacking the putative lipoprotein signal sequence) of each gene was PCR amplified using Q5 Hot Start Master Mix (New England Biolabs). Each PCR fragment, except for the one encoding ErpB, was engineered into the MCS of the pGEX4T2 expression vector (GE Healthcare Life Sciences) using BamHI and XmaI restriction sites. For *erpB*, which contains an internal BamHI restriction site, EcoRI and XmaI were used. Inserts were ligated into vector pGEX4T2 and the ligations were transformed into *E. coli* DH5α as previously described (67). Transformants were confirmed by BamHI/EcoRI and XmaI restriction digest of the plasmids, followed by gel electrophoresis on a 1% agarose gel for one hour at 75 V. Clones containing the correct insert were Sanger sequenced using an ABI 3130XL automated sequencer (Applied Biosciences). Confirmed plasmids were subsequently transformed into *E. coli* BL21(DE3) as previously described (67).

To purify GST-tagged proteins, *E. coli* BL21(DE3) cells encoding the appropriate plasmid were grown in broth culture with aeration at 37°C to an OD600nm of 0.6, then induced with 1 mM IPTG (Sigma Aldrich) with aeration at room temperature overnight. The following day, cells were lysed using an M-110S Microfluidizer (Microfluidics) and proteins were purified using glutathione chromatography according to the manufacturer’s instructions (GE Healthcare Life Sciences). To confirm the size and purity of purified recombinant protein, 25 μl of column eluate was resolved by SDS-PAGE on a 4-20% gradient polyacrylamide gel run at 75 V for 1.5 hours. Gels were then stained for 30 minutes with Coomassie blue solution [0.25% (w/v) Coomassie brilliant blue R-250, 45% (v/v) methanol, 10% (v/v) glacial acetic acid], rinsed in deionized water, and destained for two hours with destain solution [40% (v/v) methanol, 10% (v/v) glacial acetic acid]. Stained gels were imaged using a Syngene G:Box XR5 imager.

Untagged ErpQ_19-343_ was subcloned into pT7HMT by incorporating 5’ BamHI and 3’ STOP and NotI site using the pGEX4T2 construct containing ErpQ as template. Subsequent expression of BBK32-C and ErpQ_19-343_ was completed as previously described (49). All proteins used in study were assessed for purity by SDS-PAGE prior to use in assays.

### Bacterial strains, plasmids, and lipoprotein gain-of function library

*E. coli* strains DH5α and BL21(DE3) were cultured as described above. An epitope-tagged *B. burgdorferi* lipoprotein expression (“gain-of-function”) library in the high-passage, non-infectious B31-e2 background strain (6) was grown in BSK-II medium supplemented with 6% (v/v) heat-inactivated normal rabbit serum (Atlanta Biologicals) at 33°C, pH 7.6, with ambient levels of CO_2_ (68). BSK-II was supplemented with 100 μg/ml of kanamycin (Sigma Aldrich) as necessary. The gain-of-function library consists of 80 individual *B. burgdorferi* B31-e2 clones containing the low copy (approximately 10 copies per cell) pSC:LP vector, where LP represents a unique surface-exposed *B. burgdorferi* lipoprotein expressed in each of the 80 clones (Table S1). Ectopic expression of each lipoprotein-encoding gene is driven by the *B. burgdorferi flaB* constitutive promoter. The library was arrayed in multiple 96-well, sterile, flat bottom plates at a concentration of 1×10^7^ spirochetes/well (1×10^8^ per ml) and stored at -80°C in BSK-II with 20% (v/v) glycerol.

### Quantitation of binding of gain-of-function library clones to immobilized substrates

Binding of gain-of-function library clones to immobilized substrates was measured using a modification of a previously-described ELISA-based assay (51). One μg/well purified BSA (Sigma Aldrich), or human derived fibronectin (Corning) or C1 proteins (Complement Technologies) in coating buffer [0.1M sodium bicarbonate, pH 9.6] was used to coat wells of an uncoated 96-well ELISA plate (Nunc Maxisorp) at 4°C overnight. On the same day, a single 96- well plate containing the gain of function library was thawed at room temperature and centrifuged (1,250 × g, 15 min, RT) to pellet the spirochetes. The supernatant was discarded, and the cells were resuspended in 200 μl/well of BSK-II and allowed to recover under normal growth conditions. The following day, 80 μl (∼4×10^6^ cells) of culture from each well of the 96-well plate were transferred to a new plate and centrifuged (1,250 × g, 15 min, RT). The supernatant was discarded, and the cells were resuspended in 200 μl/well HBS-DB [25 mM HEPES, 105 mM sodium chloride, 1 mM manganese chloride, 1 mM magnesium chloride, 0.1% (w/v) dextrose, 0.2% (w/v) BSA, pH 7.8)]. The previously coated 96-well plate was washed three times with PBS-T [10 mM disodium phosphate, 1.8 mM monopotassium phosphate, 137 mM sodium chloride, 2.7 mM potassium chloride, 0.05% (v/v) Tween-20, pH 7.4] and was blocked with 200 μl/well of Ultrablock (BioRad) for 1.5 hours.

After discarding the blocking buffer, the plate was washed, then inoculated with 50 μl/well (∼1×10^6^ spirochetes) of the resuspended gain-of-function library. The inoculated plate was centrifuged (1,250 × g, 15 min, RT) to force the spirochetes into contact with the proteins coated on the bottoms of the wells. The plates were then incubated for one hour at room temperature, followed by three washes to remove unbound spirochetes. Bound spirochetes were affixed to the surface of the well by adding 4% formaldehyde (v/v) for 20 minutes at room temperature. After fixation, the formaldehyde was removed, and the plates were air dried on the bench top overnight.

The following day fixed spirochetes were permeabilized with 50 μl/well of ice-cold methanol for 10 minutes at -20°C. Methanol was then removed and the plates were air dried for several minutes. Wells were then blocked with 200 μl/well of 5% (w/v) non-fat dry milk in PBS-T for one hour, followed by washing. To detect spirochete binding, wells were incubated with a 1:800 dilution of a polyclonal rabbit α-Bb antibody (Abcam, ab20118) for one hour at room temperature, then washed and probed with a 1:2,000 dilution of an α-rabbit alkaline phosphatase conjugated antibody (Sigma Aldrich, cat # A3687) for one hour at room temperature. Wells were then washed and signal developed using the SigmaFast pNpp reagent (Sigma Aldrich). The assay readout (OD405nm) was taken every minute for 15 minutes using a BioTek Synergy HT plate reader and Gen5 software. Bacterial binding is expressed as the Vmean of ΔOD_405nm_, calculated by determining the slope of the [OD405nm vs. time] best fit line across the linear portion of the 15-minute kinetic assay. All experiments were repeated at least twice.

### Quantitative ELISA to assess B. burgdorferi lipoprotein binding to purified human C1

To quantitate the ability of *B. burgdorferi* lipoproteins to bind purified components of the C1 complex, we adapted a previously described quantitative ELISA-based assay (52). One μg/well of purified human C1, C1q, C1r, or C1s proteins (Complement Technologies), or BSA (Sigma-Aldrich) as a negative control, were coated onto wells of an uncoated 96-well ELISA plate (Nunc Maxisorp) overnight at 4°C in coating buffer, as described above. The next day, plates were washed three times with PBST [10 mM disodium phosphate, 1.8 mM monopotassium phosphate, 137 mM sodium chloride, 2.7 mM potassium chloride, 0.05% (v/v) Tween-20, pH 7.4] and blocked with 5% (w/v) nonfat dry milk in PBST. Plates were then washed, and 100 μl/well of four-fold dilutions of GST-tagged BBK32, ErpB, ErpQ, or BB0460 proteins, resulting in a range of concentrations from 1 μM to 240 pM, were added to the ELISA plate, which was then incubated for one hour at room temperature. Wells were washed and probed with 100 μl/well of a goat α-GST antibody (GE Healthcare Life Sciences, 27457701V) diluted 1:800 and incubated for one hour at room temperature. Wells were washed again and probed with 100 μl/well of a α-goat alkaline phosphatase conjugated antibody (Sigma Aldrich, A4187) diluted 1:2,000 and incubated for one hour at room temperature. Wells were washed a final time and the assay was developed using the SigmaFast pNpp reagent (Sigma Aldrich).

OD405nm was read every minute for 15 minutes in a BioTek Synergy HT plate reader using Gen5 software. Substrate binding is expressed as the Vmean of ΔOD405nm, calculated by determining the slope of the [OD405nm vs. time] best fit line across the linear portion of the 15-minute kinetic assay. All experiments were repeated two to four times. KD was quantified by a saturated binding parameter non-linear regression analysis performed using GraphPad Prism 6.0 software.

### Surface plasmon resonance

Binding of C1 and its sub-components to GST-ErpB and GST-ErpQ was performed at 25°C using a Biacore T200 (GE Healthcare) as previously described (Garcia 2016), with the following modifications. GST-ErpB and GST-ErpQ were amine coupled to the CMD200 (Xantec bioanalytics) at 10 μg/ml in 10 mM sodium acetate pH 4.0. Final immobilization densities shown in resonance units (RU) were 555.1 (GST-ErpB) and 451.1 RU (GST-ErpQ), and proenzyme studies were performed on 232.2 (GST-ErpB) and 485.7 (GST-ErpQ). C1s single cycle experiments had immobilization densities of 1181.3 (GST-ErpQ) and 1158.9 (ErpQ_19-343_). HBS-T-Ca^2+^ (20 mM HEPES (pH 7.3), 140 mM NaCl, 0.005% (v/v) Tween 20, 5 mM CaCl_2_) was used as the running buffer and a flowrate of 30 μl min^-1^ was used in all experiments. All analytes were buffer exchanged into running buffer prior to experimentation.

Multi cycle steady state analysis were performed as follows, C1 complex (Complement Technologies) was injected over flow cells in a two-fold concentration series: 0.59, 1.2, 2.3, 4.7, 9.4, 18.8, 37.5, 75, and 150 nM for 120 sec, followed by 180 sec dissociation. The same approach was used for proenzyme C1r, C1r enzyme, proenzyme C1s, and C1s enzyme (Complement Technologies), using a two-fold concentration series of 0.39, 0.78, 1.6, 3.1, 6.3, 13, 25, 50, 100, and 200 nM. Surfaces were then regenerated by injecting 2M NaCl for 60 sec 3 times consecutively, bringing the response to baseline. Alternatively, single cycle analysis was performed with a five-fold concentration series 0, 0.8, 4, 20, 100 nM with association times between each injection of 120 sec a final dissociation time of 600 sec. Kinetic analyses were performed on each sensorgram series using the Biacore T200 Evaluation Software 3.1 (GE Healthcare) and a 1:1 (Langmuir) binding model.

### Proteinase K treatment, conventional western and far western immunoblotting

Surface proteolysis of expressed lipoproteins was performed as previously described (34). Briefly, 1×10^8^ spirochetes were washed three times in HBS-DB and resuspended in 100 μl of HBS-DB. Spirochetes were then treated with 40 μg/ml (final concentration) of pronase (Sigma) for one hour at room temperature. Reactions were inactivated with 2 mM phenylmethanesulfonyl fluoride (PMSF) (Sigma Aldrich) and cells were lysed by boiling for ten minutes in Laemmli buffer (34). Bacterial lysates were resolved by SDS-PAGE on a 4-20% gradient polyacrylamide gel at 75 V for 1.5 hours. After electrophoresis, samples were transferred to a PVDF membrane and used for western immunoblotting.

Conventional western immunoblotting was performed by blocking the PVDF membrane with 5% (w/v) nonfat dry milk in PBS-T. Antibodies used include the CD-1 antibody (a generous gift from Jorge Benach, Stony Brook University) (1:1000 dilution) for the detection of *B. burgdorferi flaB*, and a combination of α-6×His antibody (Sigma Aldrich, H1029) and HisProbe-HRP (Thermo) for the detection of affinity-tagged lipoproteins. Following washing, immunoblots were probed with the appropriate secondary α-mouse antibody conjugated to horseradish peroxidase (Promega, W402B) at a dilution of 1:5000. Immune complexes were detected using the SuperSignal West Pico Chemiluminescent Substrate (ThermoFisher Scientific), following the manufacturer’s instructions, and imaged using a Syngene G:Box XR5 imager.

Far western immunoblotting was performed as previously described (53). Briefly, membranes were blocked in 5% (w/v) nonfat dry milk in PBS-T, followed by incubation with 2 μg/ml of purified C1q, C1r, or C1s (Complement Technologies) in protein binding buffer [20 mM Tris (pH 7.5), 0.1M sodium chloride, 1 mM EDTA, 1 mM DTT, 10% (v/v) glycerol, 0.1% (v/v) Tween-20, 5% (w/v) nonfat dry milk] overnight at 4°C. The next day, membranes were washed in PBS-T and were incubated with α-C1q (Complement Technologies, A200), α-C1r (R&D Systems, MAB1807), or α-C1s (R&D Systems, MAB2060) antibodies, following the manufacturer’s recommended dilutions for western blotting. Following washing, immunoblots were probed with the appropriate secondary α-goat (C1q) (Sigma Aldrich, A5420) or α-mouse (C1r/s) (Promega, W402B) antibodies conjugated to horseradish peroxidase, at a dilution of 1:5000. Immune complexes were detected using the SuperSignal West Pico Chemiluminescent Substrate (ThermoFisher Scientific), according to the manufacturer’s instructions, and imaged using a Syngene G:Box XR5 imager.

### Inhibition of erythrocyte hemolysis by recombinant B. burgdorferi lipoproteins

Inhibition of CP-mediated erythrocyte hemolysis by recombinant *B. burgdorferi* lipoproteins was assayed using a modified version of the previously described classical pathway hemolytic assay (28, 54). Normal human serum (Complement Technologies) was diluted to 2.3% (v/v) in CP/LP reaction buffer. GST-tagged ErpB, ErpQ, BBK32, or BB0460 proteins were serially diluted two-fold, 125 μl of each dilution was mixed with 125 μl of the diluted serum, and the mixtures were incubated at room temperature for one hour. During incubation, 5 ml of pre-opsonized sheep erythrocytes (Complement Technologies) were centrifuged (400 × g, 3 minutes, 4°C) and washed twice in CP/LP reaction buffer. After washing, erythrocytes were resuspended in 5 ml of CP/LP buffer and 40 μl of the erythrocyte suspension were added to each of the incubated serum-protein mixtures. These reactions were incubated for one hour at room temperature, gently vortexing every 15 minutes to ensure that erythrocytes remained in suspension. Following incubation, samples were centrifuged (600 × g, 3 minutes, 4°C) and 200 μl of supernatant from each sample was collected and the OD405nm was measured in a BioTek Synergy HT plate reader using Gen5 software.

### Inhibition of C3d deposition by recombinant B. burgdorferi lipoproteins

To determine the effect of recombinant *B. burgdorferi* lipoproteins on CP-mediated deposition of C3d, we adapted a previously described ELISA based assay (28, 55). 96-well ELISA plates (Nunc Maxisorp) were coated with 300 ng human IgM (CP initiator) (Athens Research & Technology) in 100 μl/well of coating buffer (see above) overnight at 4°C. The following day, the plates were washed three times with PBS-T (see above) and were blocked with 200 μl of 1% (w/v) BSA (Sigma Aldrich) in PBS-T for one hour at room temperature. Normal human serum (Complement Technologies) was diluted to 2% (v/v) in CP reaction buffer [20 mM HEPES (pH 7.3), 140 mM sodium chloride, 150 μM calcium chloride, 500 μM magnesium chloride, 0.1% (w/v) gelatin]. GST-tagged ErpB, ErpQ, BBK32, or BB0460 proteins were serially diluted two-fold and each dilution was mixed 1:1 with the above serum dilutions. 100 μl of each mixture was added to the complement initiator-treated wells. Plates were incubated in the presence of 5% CO2 for one hour at 37°C, followed by three washes with PBS-T. Following deposition, wells were blocked with 200 μl/well of 5% (w/v) nonfat dry milk in PBS-T for one hour at room temperature, followed by three washes with PBS-T. Wells were then probed with 100 μl/well of a mouse α-C3d (Abcam, ab17453) primary antibody diluted 1:500 and incubated for one hour at room temperature. Wells were washed again and probed with 100 μl/well of a α-mouse alkaline phosphatase-conjugated secondary antibody (Sigma Aldrich, A4187) diluted 1:2,000 and incubated for one hour at room temperature. Wells were washed a final time and developed using the SigmaFast pNpp reagent (Sigma Aldrich). OD405nm readings were taken every minute for 15 minutes in a BioTek Synergy HT plate reader using Gen5 software. Substrate binding is expressed as the Vmean of ΔOD_405nm_, which is calculated by determining the slope of the [OD405nm vs. time] best fit line across the linear portion of the 15- minute kinetic assay. All experiments were repeated two to four times.

### Inhibition of C4d deposition by recombinant B. burgdorferi lipoproteins

To show direct inhibition of classical pathway activation, an ELISA approach was used (28, 55). 3 ug-mL^-1^ Human IgM (Innovative Research), a classical pathway activator, in 100 mM Na_2_CO_3_/NaHCO_3_ coating buffer pH 9.6 was immobilized overnight at 37°C in high-binding polypropylene microplates (Grenier bio-one). All subsequent steps were then washed three times, 100 µl volumes, with TBS-T (50mM Tris (pH 8.0),150 mM NaCl, 0.05% (v/v) TritonX- 100). Unbound regions of the plate were then blocked with PBS-T-BSA (137mM NaCl, 2.7 mM KCl, 10mM Na_2_HPO_4_,1.8 mM KH_2_PO_4_, 1% (w/v) bovine serum albumin, and 0.05% (v/v) Tween-20) for 1 h at 37°C. Classical pathway-mediated complement activation was then induced by adding 2% final pooled Normal Human Serum (Innovative Research) and a two-fold dilution series of GST-ErpB/GST-ErpQ/GST-BB0460 or untagged proteins BBK32-C/ErpQ_19-343_, respectively, in CP Buffer (20 mM HEPES (pH 7.3), 0.1% (w/v) gelatin type A, 140 mM NaCl, 2 mM CaCl_2_,0.5 mM MgCl_2_) with incubation at 37°C for 1 hr. A 1:300 dilution anti-C4 antibody (HYB 162-02) (Santa Cruz Biotechnology) in CP Buffer incubated at 37°C for 1 hour was used to detect complement activation. A 1:3000 dilution of goat anti-mouse HRP secondary antibody (Thermo Scientific) was then used at room temperature with light rocking for 1 hour. Activation of HRP conjugated antibody was detected by room temperature 1-step Ultra TMB ELISA (Thermo Scientific) for 10 min with rocking in the dark. The reaction was then stopped with the addition of 0.16 N sulfuric acid and the absorbance measured at 450 nM on an EnSight multimode plate reader (PerkinElmer). Data were in-column normalized using cells containing serum only or no serum with buffer addition were used as 100% and 0% signal, respectively. All experiments were performed in triplicate and IC_50_ values were determined using a variable four-parameter nonlinear regression analysis using GraphPad Prism 8.1.2.

### Inhibition of C1r and C1s enzyme activity by synthetic peptide cleavage

C1r enzyme and C1s enzyme assays were performed in HBS-Ca^2+^ (20 mM HEPES (pH7.3), 140 mM NaCl, 5 mM CaCl_2_). C1r enzyme assays were completed by monitoring the autolytic activation of C1r proenzyme by adding GST-ErpB or GST-ErpQ, at a concentration of 25 μM, with 25 nM C1r proenzyme. Subsequent addition of 300 µM Z-Gly-Arg thiobenzyl (MP Biomedicals) and 100 µM 5,5′-dithiobis(2-nitrobenzoic acid) (DTNB) (TCI) just prior to measurement for a final 80 µl reaction volume (56). C1s enzyme assays were performed by adding GST-ErpB or GST-ErpQ, at a concentration of 25 μM, to 100 µM Z-L-Lys thiobenzyl and 100 µM DTNB. Just prior to measurement, 6.25 nM C1s enzyme was added for a final 80 µl reaction volume. Absorbance measurements were performed at 412 nM on a Versamax multimode plate reader (Molecular Devices) with plate reads occurring every 30 sec at 28° C for 3 hrs (C1r) and 37°C for 1 hr (C1s). Data were in-column normalized by including the C1r proenzyme or C1s enzyme with substrate as 100% signal, or just peptide and DTNB as 0%.

### Gel-Based Inhibition of C1s-Mediated C2/C4 Cleavage Assay

To demonstrate inhibition of C1s mediated cleavage of C2 or C4 a 10 µL reaction in HBS-Ca^2+^ (10 mM HEPES (pH 7.3), 140 mM NaCl, 5 mM CaCl_2_) was made by adding 6.25 nM C1s enzyme with twofold dilutions of ErpQ_19-343_ from 25,000 nM to 390 nM with subsequent addition of 1.25µL of C4 (1 mg/mL) or C2 (0.5 mg/mL) (Complement Technologies). The reaction proceeded at 37° C for 1 hour and was stopped by the addition of 5 µL Laemmli buffer followed by boiling for 5 min. 10% SDS-PAGE gels were utilized with Coomassie staining. Gel imaging was completed on a ChemiDocTM XRS+ (Bio-Rad). Gels are representative of three independent experiments.

Gel-based C2/C4 inhibition assays were subjected to further quantitative analysis with Image Lab^TM^ (Bio-Rad). Lanes and bands were manually selected and analyzed as follows. C4α’ fragments were in lane normalized to C4b band and the background adjusted ratios are shown. C2b bands were in lane corrected for total C2 (C2 + C2b + C2a). 100% cleavage was constrained to C1s + C2/C4 control. Proenzyme C1s bands were normalized to in lane C1s light chain. In the event of no detection of a band a band was placed at the appropriate analysis molecular weight. A normalized four-parameter nonparametric response was analyzed in GraphPad v8.4.

Gel-based C2/C4 inhibition assays were subjected to further quantitative analysis with Image LabTM (Bio-Rad). Lanes and bands were manually selected and analyzed as follows. C4α’ fragments were in lane normalized to C4b band and the background adjusted ratios are shown. C2b bands were in lane corrected for total C2 (C2 + C2b + C2a). 100% cleavage was constrained to C1s + C2/C4 control. Proenzyme C1s bands were normalized to in lane C1s light chain. In the event of no detection of a band a band was placed at the appropriate analysis molecular weight. A normalized four-parameter nonparametric response was analyzed in GraphPad v8.4.

### Gel-Based Inhibition of C1r-Mediated Proenzyme C1s Cleavage Assay

Enzymatic inhibition assays were performed as previously described, with the following modifications (28). A 10 µL reaction in HBS-Ca2+ (10 mM HEPES (pH 7.3), 140 mM NaCl, 5 mM CaCl_2_) was prepared by adding 1000 nM C1r to with twofold dilutions of ErpQ_19-343_ from 25,000 nM to 390 nM and finally 1 µg proenzyme C1s. The reaction was incubated at 37° C for 1 hour and was stopped by the addition of 5 µL Laemmli buffer followed by boiling for 5 min. SDS-PAGE analysis was completed as in C2/C4 cleavage assay. Gel is representative of three independent experiments.

### Classical pathway-mediated serum-killing assay

1×10^9^ B31-e2 spirochetes expressing ErpB, ErpQ, BBK32, or BB0460 were harvested in late log phase by centrifugation (4,000 × g, 15 min). Supernatant was discarded and cell pellets were washed three times in CP buffer [20 mM HEPES (pH 7.3), 140 mM NaCl, 150 μM CaCl2, 500 μM MgCl2, 0.1% gelatin]. After resuspending the pellet in 1 ml of CP buffer, the cells were split into three tubes containing 5×10^7^ spirochetes each. 4 μg of an α-*B. burgdorferi* antibody (Abcam, ab20950) was added to two tubes, while its isotype control (Abcam, ab171870) was added to the third. Cell suspensions were incubated at room temperature for 1 hour, rocking.

Following incubation, the cells were pelleted, washed three times in CP buffer, and resuspended in 625 μl of CP buffer. From one of the α-Bb antibody tubes, as well as from the isotype control tube, 1×10^7^ spirochetes (125 μl of cell suspension) was dispensed into tubes containing 125 μl of 40% normal human serum (in CP buffer), supplemented with 20 μg/ml lysozyme (Sigma, L6876), in triplicate. 125 μl of cell suspension from the other α-Bb antibody tube was dispensed into tubes containing 125 μl of 40% heat-inactivated human serum (in CP buffer), supplemented with 20 μg/ml lysozyme, in triplicate. Tubes were all mixed thoroughly by hand and incubated at 37°C, standing, for 4 hours.

Following the second incubation, the entire 250 μl of the serum-cell suspension mixture was transferred into a culture tube containing 2.25 ml of BSK-II, supplemented with the appropriate antibiotics. These cultures were allowed to grow out for 72 hours in normal growth conditions. After 72 hours of growth, the cultures were counted in duplicate by dark field microscopy. Samples were normalized to triplicate counts from the heat-inactivated human serum samples.

## Acknowledgments

Research reported in this publication was supported by the National Institute of Allergy and Infectious Diseases of the National Institutes of Health under award number R01AI146930 (B.L.G), R01AI121401 (JML), K12GM133314 (JDQ). We thank Yi-Pin Lin and Jon Skare for invaluable discussion, and Brian Stevenson for providing anti-Erp antibodies.

## Author Contributions

- Designed research: MJP, BW, PK, BLG, JML
- Performed research: MJP, BW, RJG, PK, EG
- Contributed new reagents or analytic tools; AD, WRZ
- Analyzed data: MJP, BW, MSO, RJG, PK, BLG, JML
- Wrote the paper: MJP, MSO, JDQ, BW, RJG, BLG, JML

