## Supplemental Figures for "Lipoproteome screening of the Lyme disease agent identifies novel inhibitors of antibody-mediated complement killing"

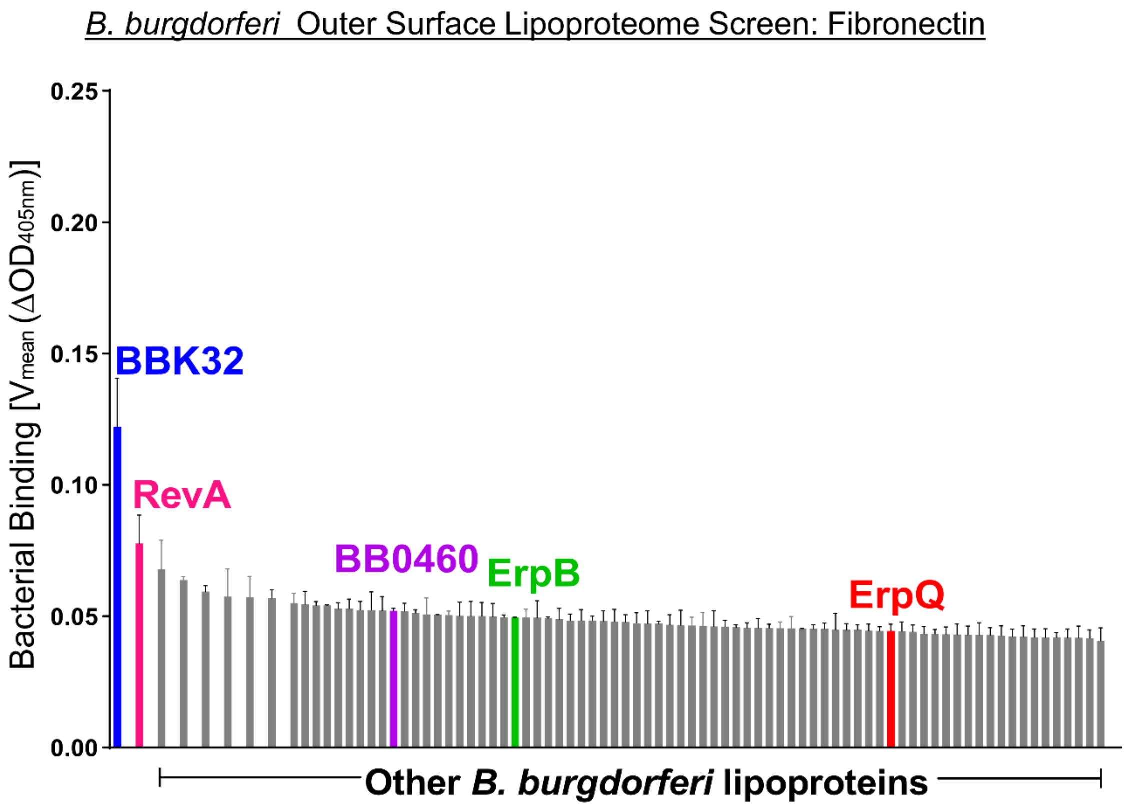


**Figure S1. Validation of the Outer Surface Lipoproteome Screening Assay Using Human Fibronectin.** A 96-well plate was coated with human fibronectin. Abbreviated names of the overproduced lipoproteins are shown on the x-axis and relative bacterial binding, reported as the change in $OD_{405nm}$ over time, is reported on the y-axis. The data are sorted from left to right in order of efficiency of binding. Error bars indicate SEM. The two *B. burgdorferi* lipoproteins that have previously been identified as capable of binding directly to human fibronectin, BBK32 (blue) and RevA (pink), were also the two highest binding lipoproteins in the screen. Proteins used in this study for C1-binding are marked: ErpQ (red), ErpB (green), BB0460 (purple).

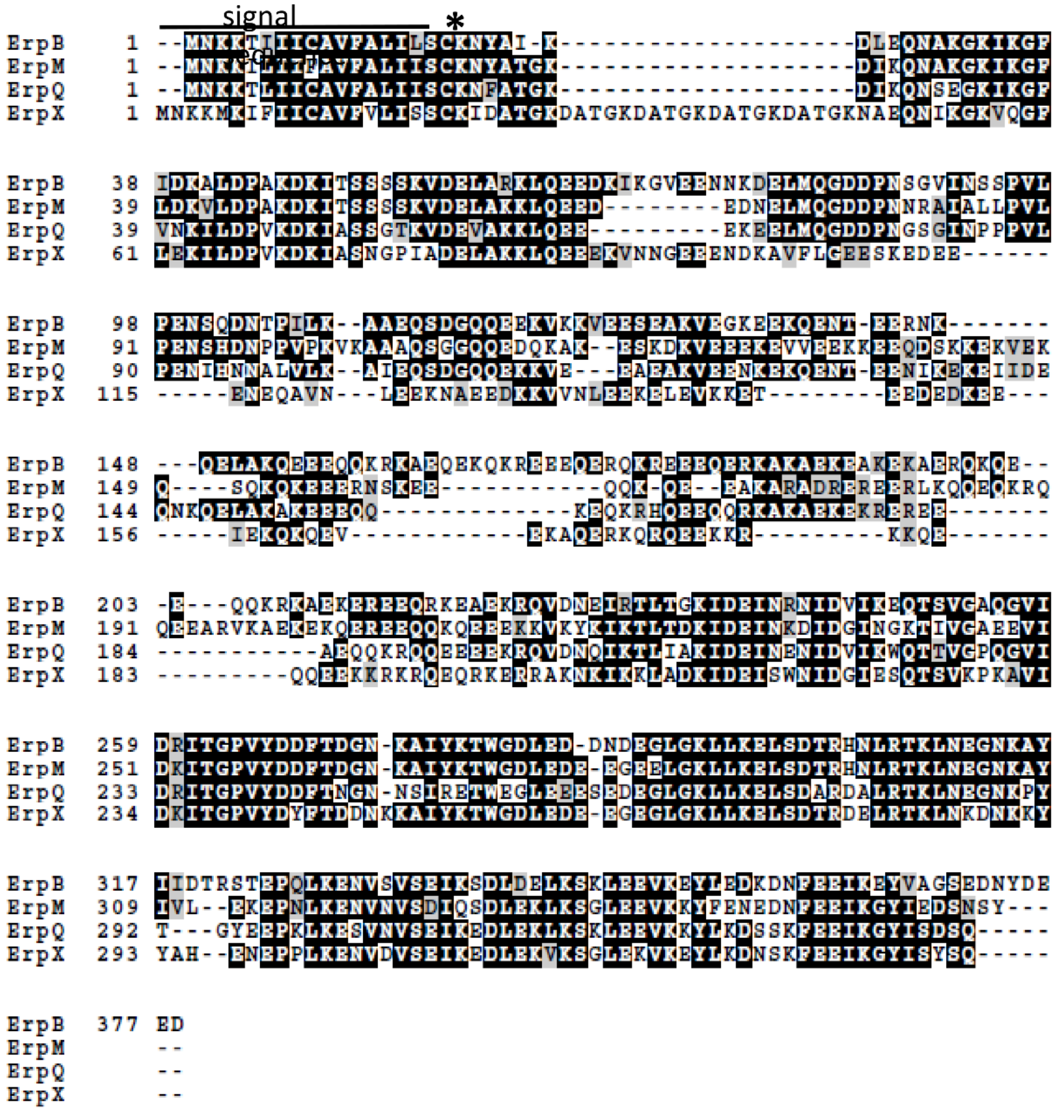


**Figure S2. BOXSHADE alignment of Elp proteins. Asterisk indicates the site of lipoprotein acylation.**

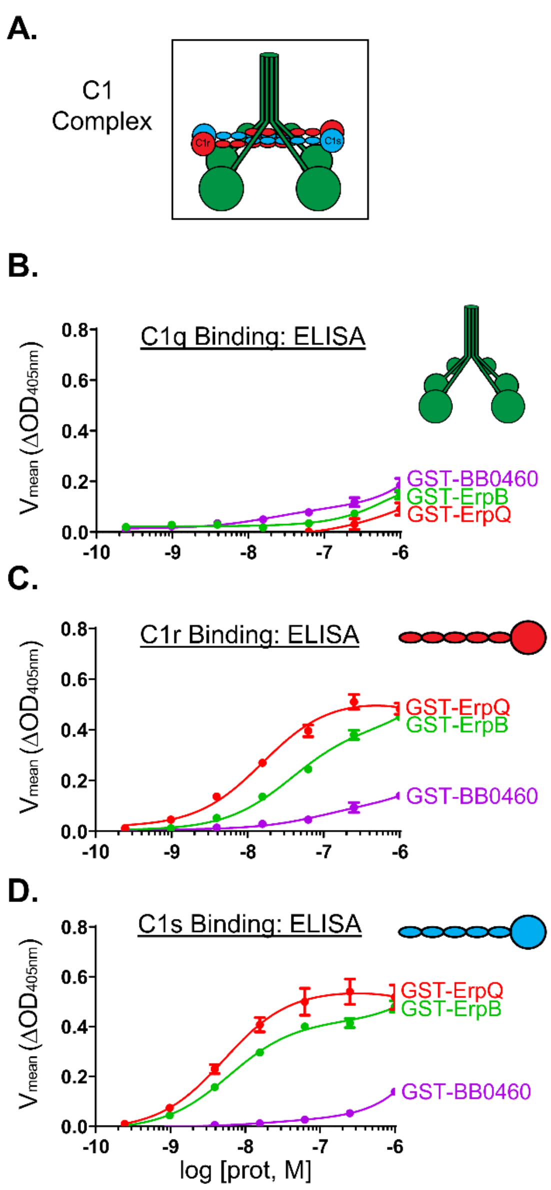


**Figure S3. ELISA-type Binding Assay. A)** Model of C1 complex subcomponent structure. C1q (top, green), C1r (middle, red), and C1s (bottom, cyan). A four-fold dilution series of the indicated purified GST-fusion proteins were applied to wells coated with (B) C1q, (C) C1r enzyme, (D) or C1s enzyme. Substrate binding is reported as the change in $OD_{405\ nm}$ over time. Error bars indicate SEM. Affinity analysis was performed on Prism GraphPad software, using a non-linear regression analysis fitting procedure.

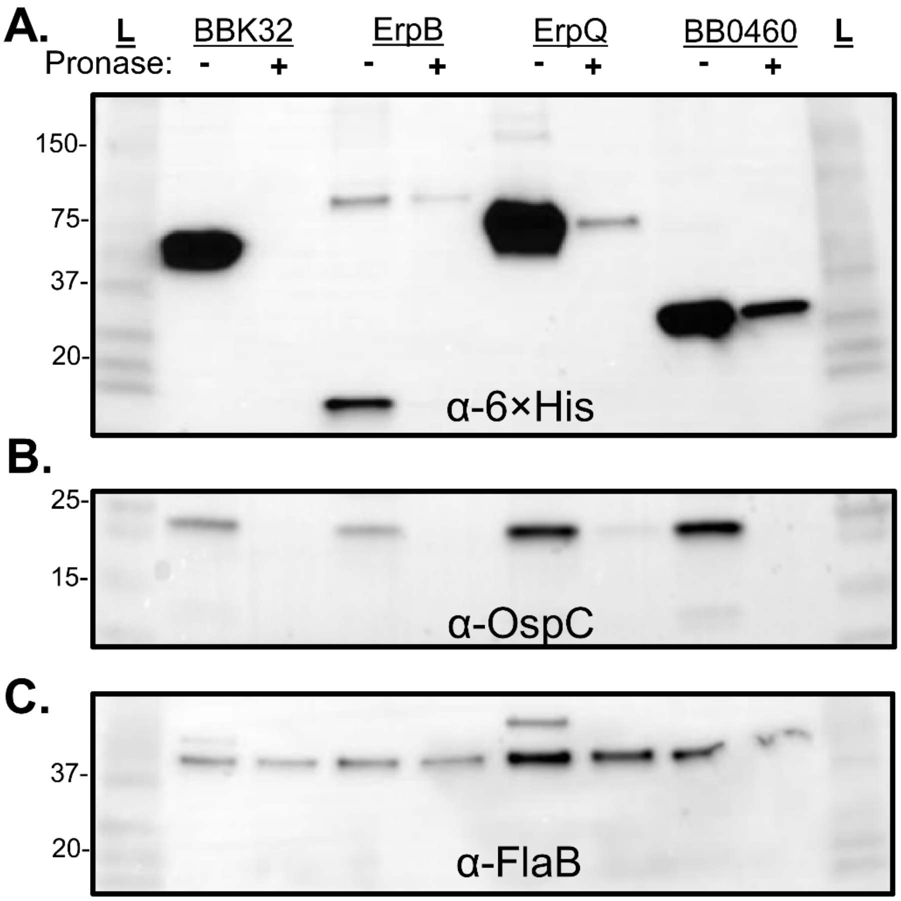


**Figure S4. Validation of apparent molecular weight and localization of epitope tagged BBK32, ErpB, ErpQ, and BB0460 expressed in *B. burgdorferi* strain B31-e2.** $1\times10^7$ spirochetes +/-pronase treatment were lysed, resolved by SDS-PAGE, and transferred to a PVDF membrane. **A)** An α-6×His antibody, combined with HRP-conjugated $Ni^{2+}$ beads, were used to determine "prey" levels in both treatment conditions. All observed bands correspond to the reported sizes of each protein. **B)** Antibodies to OspC or **C)** FlaB were used to assess pronase accessibility to the bacterial surface and periplasm, respectively.

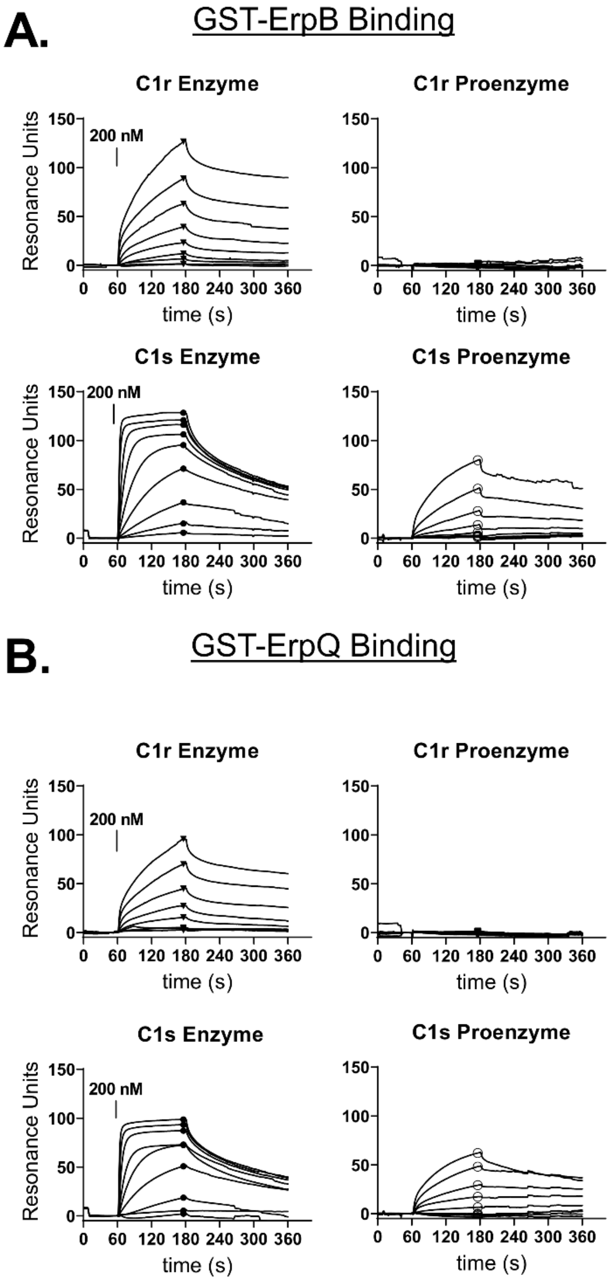


**Figure S5. SPR sensorgrams for GST-ErpB and GST-ErpQ binding to C1r and C1s proenzyme and enzyme forms.** A) GST-ErpB or B) GST-ErpQ raw sensorgrams are shown for C1r ezyme (top left), C1r proenzyme (top right), C1s enzyme (top left) and C1s proenzyme (bottom right). Symbols indicate the region of the sensorgram treated as steady-state signal for the fits shown in **Fig. 2 C, D**.

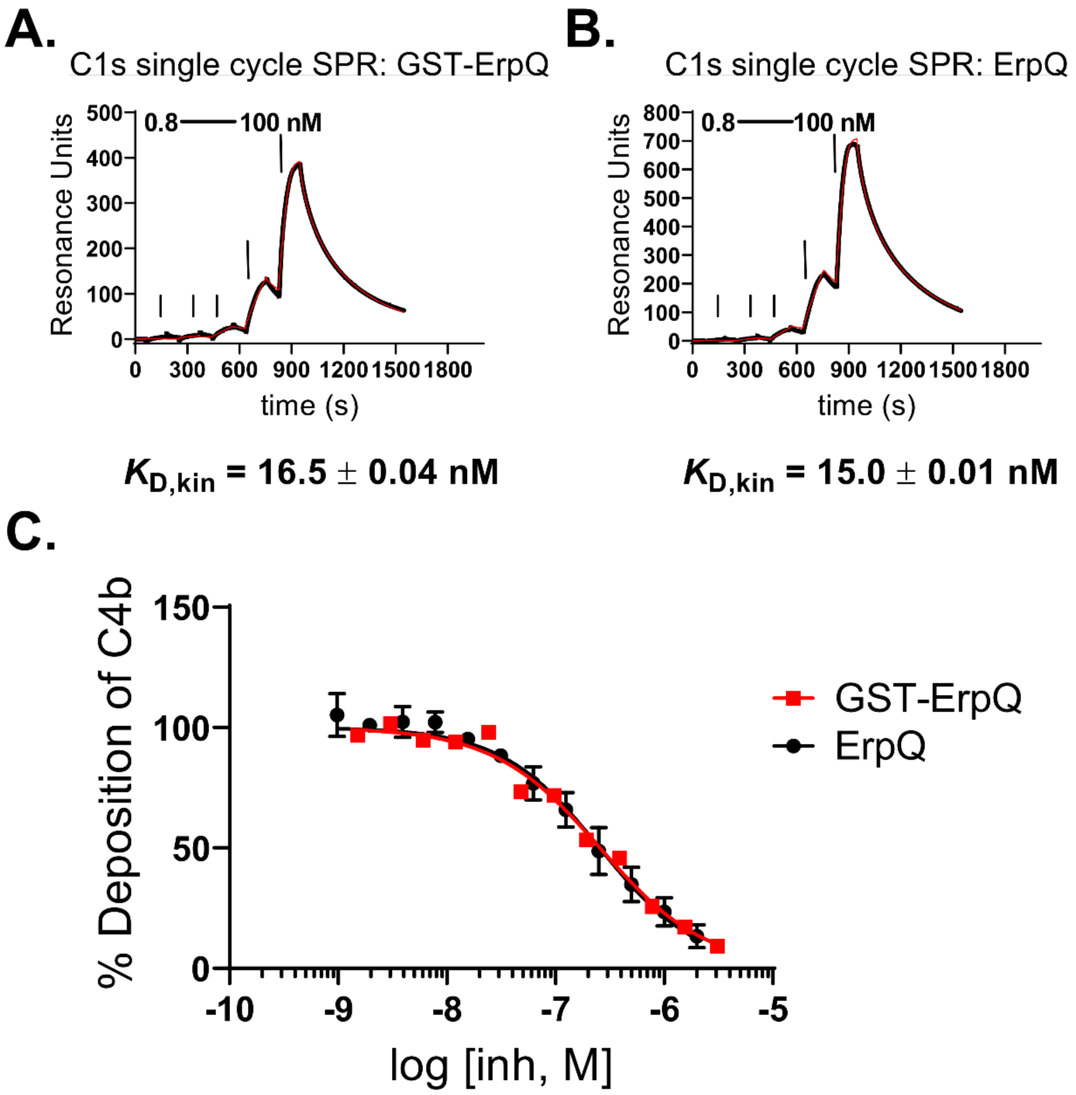


**Figure S6. Validation of ErpQ Lacking a GST-affinity Tag. A-B)** The interaction of C1s enzyme with GST-ErpQ and ErpQ lacking a GST tag was measured by SPR. A single cycle kinetic analysis was used for determination of $K_D$ values (i.e. $K_{D,kin}$). A five-fold injection series (0, 0.8, 4, 20, and 100 nM) was repeated in triplicate. **C)** A classical pathway-specific ELISA-based complement assay using C4b detecting was performed for ErpQ lacking a GST. The data for GST-ErpQ presented in Fig. 4A are replotted here for comparisons sake. Each ErpQ concentration series was repeated in triplicate.

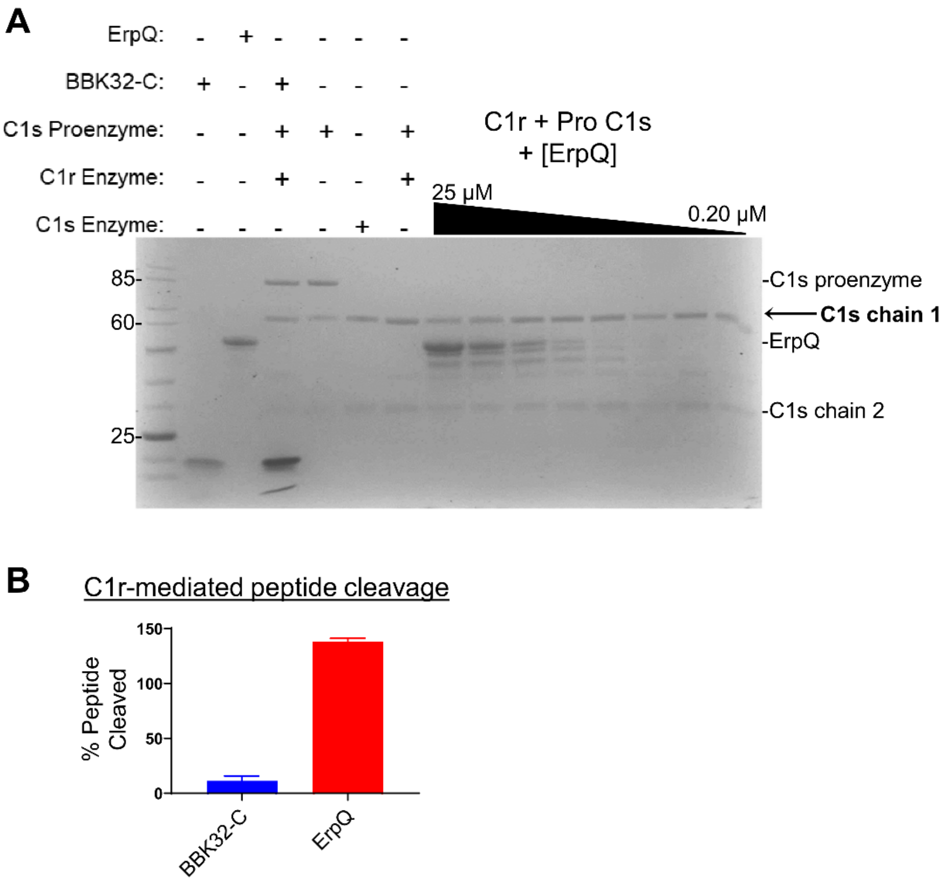


**Figure S7. C1r enzyme Inhibition Assays. A)** C1r enzymatic cleavage of active site analog Z-Gly-Arg-sBzl was assayed with DTNB (Ellman's reagent) in the presence of inhibitor (25 μM) at 25°C for 1hr. Absorbance was read at 412nm and signals were normalized to enzyme only wells. BBK32-C was used a positive control for inhibition of C1r. **B)** The ability of ErpQ to block the C1r enzyme-mediated cleavage of C1s proenzyme was assayed by monitored by SDS-PAGE. Upon cleavage by C1r, C1s proenzyme is converted to two polypeptide chains of ~58 (chain 1) and ~28 kDa (chain 2). Lane 1: Ladder, Lanes (2-7) various combinations of proteins in the assay denoting addition (+) or absence (-) of reagent. Lanes (8-15) concentrations of ErpQ up to 25 μM failed to prevent the cleavage of C1s proenzyme by C1r enzyme. BBK32-C was used as a positive control for C1r inhibition.

**Table S1. Plasmids used in this study.**

| Name | Vector | Size (kb) | Insert | Size (bp) | Reference |
|---|---|---|---|---|---|
| pSC1116 | pSC:LP | 6.3 | *bbk32* | 1081 | (6) |
| pSC1070 | pSC:LP | 6.3 | *erpB* | 1153 | (6) |
| pSC1085 | pSC:LP | 6.3 | *erpQ* | 1048 | (6) |
| pSC1126 | pSC:LP | 6.3 | *erpX* | 1054 | (6) |
| pSC1079 | pSC:LP | 6.3 | *erpM* | 1108 | (6) |
| pSC1028 | pSC:LP | 6.3 | *bb_0460* | 730 | (6) |
| pMP15 | pGEX4T2 | 5 | *bb_0460* | 660 | This study |
| pMP16 | pGEX4T2 | 5 | *bbk32* | 1002 | This study |
| pMP17 | pGEX4T2 | 5 | *erpQ* | 975 | This study |
| pMP18 | pGEX4T2 | 5 | *erpB* | 1080 | This study |
| pErpQ | pT7HMT | 5.4 | *erpQ* | 975 | This study |
| pBBK32-C | pT7HMT | 5.4 | *bbk32 206-348* | 426 | Garcia 2016 |
| pSC1037 | pSC:LP | 6.3 | *bb_0158* | 773 | (6) |
| pSC1048 | pSC:LP | 6.3 | *bb_0171* | 604 | (6) |
| pSC1047 | pSC:LP | 6.3 | *bb_0213* | 670 | (6) |
| pSC1016 | pSC:LP | 6.3 | *bb_0689* | 484 | (6) |
| pSC1039 | pSC:LP | 6.3 | *bb_0758* | 799 | (6) |
| pSC1042 | pSC:LP | 6.3 | *bb_0823* | 388 | (6) |
| pSC1059 | pSC:LP | 6.3 | *bb_A04* | 865 | (6) |
| pSC1061 | pSC:LP | 6.3 | *bb_A07* | 505 | (6) |
| pSC1068 | pSC:LP | 6.3 | *bb_A14* | 382 | (6) |
| pSC1000 | pSC:LP | 6.3 | *ospA* | 958 | (6) |
| pSC1009 | pSC:LP | 6.3 | *ospB* | 907 | (6) |
| pSC1008 | pSC:LP | 6.3 | *dbpA* | 592 | (6) |
| pSC1065 | pSC:LP | 6.3 | *bb_A32* | 211 | (6) |
| pSC1054 | pSC:LP | 6.3 | *bb_A33* | 556 | (6) |
| pSC1063 | pSC:LP | 6.3 | *bb_A36* | 658 | (6) |
| pSC1053 | pSC:LP | 6.3 | *bb_A57* | 1261 | (6) |
| pSC1057 | pSC:LP | 6.3 | *bb_A59* | 256 | (6) |
| pSC1060 | pSC:LP | 6.3 | *bb_A64* | 940 | (6) |
| pSC1064 | pSC:LP | 6.3 | *bb_A65* | 865 | (6) |
| pSC1062 | pSC:LP | 6.3 | *cspA* | 772 | (6) |
| pSC1058 | pSC:LP | 6.3 | *bb_A69* | 808 | (6) |
| pSC1014 | pSC:LP | 6.3 | *bb_B08* | 646 | (6) |
| pSC1002 | pSC:LP | 6.3 | *ospC* | 649 | (6) |

| | | | | | |
|---|---|---|---|---|---|
| pSC1025 | pSC:LP | 6.3 | *bb_B25* | 529 | (6) |
| pSC1087 | pSC:LP | 6.3 | *bb_C10* | 544 | (6) |
| pSC1088 | pSC:LP | 6.3 | *bb_D10* | 595 | (6) |
| pSC1091 | pSC:LP | 6.3 | *bb_E04* | 157 | (6) |
| pSC1089 | pSC:LP | 6.3 | *bb_E08* | 163 | (6) |
| pSC1090 | pSC:LP | 6.3 | *bb_E31* | 739 | (6) |
| pSC1093 | pSC:LP | 6.3 | *bb_F01* | 1015 | (6) |
| pSC1092 | pSC:LP | 6.3 | *bb_F20* | 310 | (6) |
| pSC1095 | pSC:LP | 6.3 | *bb_G01* | 910 | (6) |
| pSC1099 | pSC:LP | 6.3 | *bb_H01* | 211 | (6) |
| pSC1098 | pSC:LP | 6.3 | *cspZ* | 727 | (6) |
| pSC1096 | pSC:LP | 6.3 | *bb_H18* | 1132 | (6) |
| pSC1097 | pSC:LP | 6.3 | *bb_H32* | 766 | (6) |
| pSC1100 | pSC:LP | 6.3 | *bb_H37* | 955 | (6) |
| pSC1107 | pSC:LP | 6.3 | *bb_I14* | 130 | (6) |
| pSC1102 | pSC:LP | 6.3 | *vraA* | 1372 | (6) |
| pSC1101 | pSC:LP | 6.3 | *bb_I28* | 589 | (6) |
| pSC1108 | pSC:LP | 6.3 | *bb_I29* | 682 | (6) |
| pSC1104 | pSC:LP | 6.3 | *bb_I36* | 853 | (6) |
| pSC1105 | pSC:LP | 6.3 | *bb_I38* | 853 | (6) |
| pSC1106 | pSC:LP | 6.3 | *bb_I39* | 883 | (6) |
| pSC1103 | pSC:LP | 6.3 | *bb_I42* | 571 | (6) |
| pSC1121 | pSC:LP | 6.3 | *bb_J01* | 202 | (6) |
| pSC1011 | pSC:LP | 6.3 | *ospD* | 790 | (6) |
| pSC1119 | pSC:LP | 6.3 | *bb_J34* | 1087 | (6) |
| pSC1118 | pSC:LP | 6.3 | *bb_J36* | 1075 | (6) |
| pSC1120 | pSC:LP | 6.3 | *bb_J41* | 883 | (6) |
| pSC1117 | pSC:LP | 6.3 | *bb_K01* | 910 | (6) |
| pSC1111 | pSC:LP | 6.3 | *bb_K07* | 769 | (6) |
| pSC1110 | pSC:LP | 6.3 | *bb_K12* | 715 | (6) |
| pSC1113 | pSC:LP | 6.3 | *bb_K19* | 652 | (6) |
| pSC1112 | pSC:LP | 6.3 | *bb_K48* | 883 | (6) |
| pSC1109 | pSC:LP | 6.3 | *bb_K50* | 1015 | (6) |
| pSC1115 | pSC:LP | 6.3 | *bb_K52* | 865 | (6) |
| pSC1114 | pSC:LP | 6.3 | *bb_K53* | 571 | (6) |
| pSC1083 | pSC:LP | 6.3 | *mlpH* | 463 | (6) |
| pSC1081 | pSC:LP | 6.3 | *erpN* | 541 | (6) |
| pSC1076 | pSC:LP | 6.3 | *mlpF* | 466 | (6) |
| pSC1075 | pSC:LP | 6.3 | *erpK* | 787 | (6) |

| | | | | | |
|---|---|---|---|---|---|
| pSC1086 | pSC:LP | 6.3 | *mlpI* | 445 | (6) |
| pSC1084 | pSC:LP | 6.3 | *erpP* | 568 | (6) |
| pSC1071 | pSC:LP | 6.3 | *revA* | 490 | (6) |
| pSC1069 | pSC:LP | 6.3 | *mlpA* | 463 | (6) |
| pSC1010 | pSC:LP | 6.3 | *erpA* | 529 | (6) |
| pSC1125 | pSC:LP | 6.3 | *bb_Q03* | 568 | (6) |
| pSC1124 | pSC:LP | 6.3 | *bb_Q05* | 760 | (6) |
| pSC1127 | pSC:LP | 6.3 | *mlpJ* | 628 | (6) |
| pSC1128 | pSC:LP | 6.3 | *bb_Q89* | 211 | (6) |
| pSC1074 | pSC:LP | 6.3 | *mlpD* | 439 | (6) |
| pSC1073 | pSC:LP | 6.3 | *bb_R40* | 121 | (6) |
| pSC1012 | pSC:LP | 6.3 | *bb_R42* | 691 | (6) |
| pSC1072 | pSC:LP | 6.3 | *mlpC* | 463 | (6) |
| pSC1013 | pSC:LP | 6.3 | *erpG* | 607 | (6) |

**Table S2. Fn- and C1-binding by *Bb* strain B31-e2 that ectopically produce *Bb* surface lipoproteins**

| Gene locus | Location | Gene name | Fn-binding [a] | C1-binding[b] |
|---|---|---|---|---|
| *bb_0460*[d] | chromosome | | 0.0521 | 0.0488 |
| *bb_0158* | chromosome | | 0.0519 | 0.0576 |
| *bb_0171* | chromosome | | 0.0501 | 0.0553 |
| *bb_0213* | chromosome | | 0.0501 | 0.0522 |
| *bb_0689* | chromosome | | 0.0495 | 0.0511 |
| *bb_0758* | chromosome | | 0.0501 | 0.0486 |
| *bb_0823* | chromosome | | 0.0521 | 0.0484 |
| *bb_A04* | lp54 | | 0.0550 | 0.0504 |
| *bb_A07* | lp54 | *chpAI* | 0.0546 | 0.0492 |
| *bb_A14* | lp54 | | 0.0529 | 0.0521 |
| *bb_A15* | lp54 | *ospA* | 0.0570 | 0.0533 |
| *bb_A16* | lp54 | *ospB* | 0.0541 | 0.0510 |
| *bb_A24* | lp54 | *dbpA* | 0.0594 | 0.0504 |
| *bb_A32* | lp54 | | 0.0499 | 0.0520 |
| *bb_A33* | lp54 | | 0.0460 | 0.0481 |
| *bb_A36* | lp54 | | 0.0429 | 0.0469 |
| *bb_A57* | lp54 | | 0.0456 | 0.0454 |
| *bb_A59* | lp54 | | 0.0433 | 0.0441 |
| *bb_A64* | lp54 | | 0.0449 | 0.0450 |
| *bb_A65* | lp54 | | 0.0464 | 0.0443 |
| *bb_A68* | lp54 | *cspA* | 0.0455 | 0.0422 |
| *bb_A69* | lp54 | | 0.0452 | 0.0483 |
| *bb_B08* | cp26 | | 0.0529 | 0.0500 |
| *bb_B19* | cp26 | *ospC* | 0.0493 | 0.0490 |
| *bb_B25* | cp26 | | 0.0523 | 0.0518 |
| *bb_C10* | cp9 | *revB* | 0.0449 | 0.0478 |
| *bb_D10* | lp17 | | 0.0440 | 0.0451 |
| *bb_E04* | lp25 | | 0.0416 | 0.0415 |
| *bb_E08* | lp25 | | 0.0407 | 0.0405 |
| *bb_E31* | lp25 | | 0.0479 | 0.0562 |
| *bb_F01* | lp28-1 | *erpD* | 0.0420 | 0.0427 |
| *bb_F20* | lp28-1 | | 0.0419 | 0.0399 |

| | | | | |
|---|---|---|---|---|
| *bb_G01* | lp28-2 | | 0.0454 | 0.0428 |
| *bb_H01* | lp28-3 | | 0.0453 | 0.0423 |
| *bb_H06* | lp28-3 | *cspZ* | 0.0444 | 0.0427 |
| *bb_H18* | lp28-3 | | 0.0505 | 0.0459 |
| *bb_H32* | lp28-3 | | 0.0506 | 0.0506 |
| *bb_H37* | lp28-3 | | 0.0507 | 0.0508 |
| *bb_I14* | lp28-4 | | 0.0420 | 0.0459 |
| *bb_I16* | lp28-4 | *vraA* | 0.0418 | 0.0407 |
| *bb_I28* | lp28-4 | | 0.0463 | 0.0425 |
| *bb_I29* | lp28-4 | | 0.0512 | 0.0491 |
| *bb_I36* | lp28-4 | | 0.0496 | 0.0476 |
| *bb_I38* | lp28-4 | | 0.0456 | 0.0428 |
| *bb_I39* | lp28-4 | | 0.0575 | 0.0510 |
| *bb_I42* | lp28-4 | | 0.0443 | 0.0403 |
| *bb_J01* | lp38 | | 0.0430 | 0.0408 |
| *bb_J09* | lp38 | *ospD* | 0.0428 | 0.0402 |
| *bb_J34* | lp38 | | 0.0420 | 0.0408 |
| *bb_J36* | lp38 | | 0.0680 | 0.0552 |
| *bb_J41* | lp38 | | 0.0573 | 0.0476 |
| *bb_K01* | lp36 | | 0.0495 | 0.0432 |
| *bb_K07* | lp36 | | 0.0472 | 0.0425 |
| *bb_K12* | lp36 | | 0.0433 | 0.0412 |
| *bb_K19* | lp36 | | 0.0431 | 0.0431 |
| *bb_K32* | lp36 | *bbk32* | 0.1221 | 0.1425 |
| *bb_K48* | lp36 | | 0.0454 | 0.0415 |
| *bb_K50* | lp36 | | 0.0638 | 0.0485 |
| *bb_K52* | lp36 | | 0.0426 | 0.0421 |
| *bb_K53* | lp36 | | 0.0430 | 0.0436 |
| *bb_L28* | cp32-8 | *mlpH* | 0.0478 | 0.0468 |
| *bb_L39* | cp32-8 | *erpN* | 0.0489 | 0.0481 |
| *bb_M28* | cp32-6 | *mlpF* | 0.0445 | 0.0479 |
| *bb_M38* | cp32-6 | *erpK* | 0.0422 | 0.0431 |
| *bb_N28* | cp32-9 | *mlpI* | 0.0449 | 0.0438 |
| *bb_N38* | cp32-9 | *erpP* | 0.0423 | 0.0438 |
| *bb_N39* | cp32-9 | *erpQ* | 0.0444 | 0.2230 |
| *bb_P27* | cp32-1 | *revA* | 0.0777 | 0.0444 |
| *bb_P28* | cp32-1 | *mlpA* | 0.0453 | 0.0440 |

| | | | | |
|---|---|---|---|---|
| *bb_P38* | cp32-1 | *erpA* | 0.0458 | 0.0439 |
| *bb_P39* | cp32-1 | *erpB* | 0.0495 | 0.2064 |
| *bb_Q03* | lp56 | | 0.0484 | 0.0479 |
| *bb_Q05* | lp56 | | 0.0541 | 0.0515 |
| *bb_Q35* | lp56 | *mlpJ* | 0.0523 | 0.0540 |
| *bb_Q47* | lp56 | *erpX* | 0.0478 | 0.0703 |
| *bb_Q89* | lp56 | | 0.0468 | 0.0487 |
| *bb_R28* | cp32-4 | *mlpD* | 0.0474 | 0.0495 |
| *bb_R40* | cp32-4 | *erpH* | 0.0461 | 0.0496 |
| *bb_R42* | cp32-4 | *erpY* | 0.0482 | 0.0483 |
| *bb_S30* | cp32-3 | *mlpC* | 0.0466 | 0.0514 |
| *bb_S41* | cp32-3 | *erpG* | 0.0484 | 0.0479 |

[a]Expressed as change in $OD_{405nm}$ over time in ELISA-based quantitation of binding to immobilized Fn, as described in Fig. S1 legend and Methods.

[b]Expressed as change in $OD_{405nm}$ over time in ELISA-based quantitation of binding to immobilized C1, as described in Fig. 1 legend and Methods.

[c]Not applicable.

[d]Encodes a periplasmic lipoprotein and was used as a negative control.

**Table S3. Degree of identity and similarity among *B. burgdorferi* B31 Elp proteins.**

| | ErpB | ErpM | ErpO | ErpQ | ErpX |
|---|---|---|---|---|---|
| ErpB | | 75.7% | 100% | 73.1% | 59.4% |
| ErpM | 58.9% | | 75.7% | 70.1% | 59.0% |
| ErpO | 100% | 58.9% | | 73.1% | 59.4% |
| ErpQ | 58.0% | 54.9% | 58.0% | | 63.5% |
| ErpX | 43.6% | 45.4% | 43.6% | 47.4% | |

% Similarity (above diagonal)

% Identity (below diagonal)

**Table S4. Primers used in this study.**

| Name | Primer Sequence | Notes |
|---|---|---|
| prMP164 | gctagGGATCCTCTAAATCAGTCTCAAGTA | Forward primer to amplify the *bb_0460* gene |
| prMP165 | gctagCCCGGGAGACTGAGAGTG | Reverse primer to amplify the *bb_0460* gene |
| prMP177 | gctagGGATCCGATTTATTCATAAGATATG | Forward primer to amplify the *bbk32* gene |
| prMP178 | gctagCCCGGGGTACCAAACGC | Reverse primer to amplify the *bbk32* gene |
| prMP179 | gctagGGATCCAAGAATTTTGCAACT | Forward primer to amplify the *erpQ* gene |
| prMP180 | gctagCCCGGGCTGACTGTCA | Reverse primer to amplify the *erpQ* gene |
| prMP181 | gctagGAATTCCTAAGAATTATGCAATTAAAG | Forward primer to amplify the *erpB* gene |
| prMP182 | gctagCCCGGGATCTTCTTCATCATA | Reverse primer to amplify the *erpB* gene |
| ErpQ$_{19-343}$F | ggatccAAGAATTTTGCAACTGGTAA | Forward primer to amplify the *erpQ* gene for pT7HMT |
| ErpQ$_{19-343}$R | gcggccgcTTACTGACTGTCACTGATGTATC | Reverse primer to amplify the *erpQ* gene for pT7HMT |
